# Spinocerebellar ataxia type 11-associated alleles of *Ttbk2* dominantly interfere with ciliogenesis and cilium stability

**DOI:** 10.1101/274266

**Authors:** Emily Bowie, Ryan Norris, Kathryn V. Anderson, Sarah C. Goetz

## Abstract

Spinocerebellar ataxia type 11 (SCA11) is a rare, dominantly inherited human ataxia characterized by atrophy of Purkinje neurons in the cerebellum. SCA11 is caused by mutations in the gene encoding the Serine/Threonine kinase Tau tubulin kinase 2 (TTBK2) that result in premature truncations of the protein. We previously showed that TTBK2 is a key regulator of the assembly of primary cilia *in vivo*. However, the mechanisms by which the SCA11-associated mutations disrupt TTBK2 function, and whether they interfere with ciliogenesis were unknown. In this work, we present evidence that SCA11-associated mutations are dominant negative alleles and that the resulting truncated protein (TTBK2^SCA11^) interferes with the function of full length TTBK2 in mediating ciliogenesis. A *Ttbk2* allelic series revealed that upon partial reduction of full length TTBK2 function, TTBK2^SCA11^ can interfere with the activity of the residual wild-type protein to decrease cilia number and interrupt cilia-dependent Sonic hedgehog (SHH) signaling. Our studies have also revealed new functions for TTBK2 after cilia initiation in the control of cilia length, trafficking of a subset of SHH pathway components, including Smoothened (SMO), and cilia stability. These studies provide a molecular foundation to understand the cellular and molecular pathogenesis of human SCA11, and help account for the link between ciliary dysfunction and neurodegenerative diseases.

**Author Summary:** Defects in primary cilia structure and function are linked to a number of recessive genetic disorders, now collectively referred to as ciliopathies. Most of the characteristics of these disorders arise from disruptions to embryonic development, with the requirements for primary cilia in adult tissues being less well-defined. We previously showed that a kinase associated with an adult-onset neurodegenerative condition is required for cilium assembly and ciliary signaling during development. Here, we show that the human disease-associated mutations act as mild dominant negatives, interfering with the function of the full length protein in cilia formation and ciliary signaling.

## Introduction

Primary cilia play a critical role in many aspects of embryonic development. Cilia are important for the development of the brain and central nervous system, which accounts for the structural brain defects, cognitive impairments and other neurological disorders that are characteristic of many human ciliopathies[1–3]. Cilia are present on a wide variety of neurons and astroglia within the adult brain, although the specific requirements for these organelles in the function of the adult brain are not well understood.

In prior work, we identified a Serine/Threonine kinase, Tau tubulin kinase 2 (TTBK2), that is essential for initiating the assembly of primary cilia in the embryo[4]. TTBK2 is a 1244 amino acid protein (1243 amino acids in mouse) that was initially purified from bovine brain tissue as a microtubule-associated protein[5–7]. The protein is comprised of a kinase domain (AA 21-284), and a long C-terminus that is important for targeting TTBK2 to the mother centriole[4], as well as mediating its interactions with end-binding proteins at the microtubule +tips[8], and likely for additional regulation of TTBK2 function. TTBK2 was initially shown to phosphorylate the microtubule-associated proteins TAU and MAP2 in addition to β-Tubulin *in vitro[6]*, and more recent evidence suggests that TTBK2 can phosphorylate the centriolar distal appendage protein CEP164[9], as well as the atypical kinesin KIF2A at the microtubule +tips[10].

In addition to the critical requirement for TTBK2 in ciliogenesis, particular dominant mutations that disrupt TTBK2 cause a hereditary ataxia, spinocerebellar ataxia type 11 (SCA11)[11]. Like other subtypes of SCA, SCA11 is a progressive neurodegenerative condition predominantly affecting the cerebellum. At the cellular level, SCA11 is characterized by cerebellar atrophy resulting from a degeneration of Purkinje cells (PCs) of the cerebellum. However, the molecular basis underlying this pathology as well as for the dominant mode of SCA11 inheritance remain unknown. Three different heterozygous familial *SCA11*-associated mutations in *TTBK2* that cause late-onset ataxia are associated with insertions or deletions of one or two bases that cause frame shifts and produce similar truncations of TTBK2 protein C-terminal to the kinase domain, at approximately AA 450[11,12]. A fourth mutation in *TTBK2* that causes an earlier-onset disease truncates the protein at AA 402[13].

Because TTBK2 is essential for the biogenesis of primary cilia, which are in turn essential for the development of the nervous system, we hypothesized that the SCA11-associated mutations disrupt the function of TTBK2 in cilia formation. In previous structure-function experiments, we tested the ability of truncations of TTBK2 to restore cilia in *Ttbk2* null mutant cells and found that those corresponding with the SCA11-associated mutations were unable to rescue cilia formation. We also found that, when over-expressed in Wild-Type (WT) cells by retroviral transduction, these truncations also partially suppress cilia formation[4], suggesting the SCA11-associated truncations may act by interfering with the activity of the wild-type gene product.

In the present study, we examined phenotypes of mice with *SCA11*-like truncating mutations knocked into the endogenous *Ttbk2* locus. We found that, with respect to cilia, *Ttbk2^sca11^* homozygotes are indistinguishable from the null allele. Specifically, in these mutants, cilia initiation fails and cilia-dependent SHH signaling is blocked. Using a series of *Ttbk2* alleles, we showed that SCA11-associated truncated proteins dominantly interfere with the function of full-length (wild-type) TTBK2 in cilium assembly. In addition, these allelic combinations have uncovered a previously unappreciated function for TTBK2 in the regulation of cilia stability. TTBK2 localizes to the mother centriole prior to cilia formation and remains at the transition zone of the cilium following completion of assembly. In this study, we present evidence from hypomorphic allelic combinations that TTBK2 also acts after cilium initiation to regulate cilium stability, in part by countering a cilium disassembly pathway.

## Results

### Embryos homozygous for a familial SCA11-associated mutation in *Ttbk2* phenocopy *Ttbk2* null embryos

To examine the effects of SCA11-associated TTBK2 truncations on the function of the protein in mediating cilia formation and function, we used an allele of *Ttbk2* in which a mutation precisely recapitulating one of the human SCA11-causing mutations was knocked into the mouse *Ttbk2* genomic locus[14]. *Ttbk2^sca11/sca11^* homozygous embryos were previously reported to die by E11, but their developmental and cellular phenotypes were not described[15]. We found that E10.5 *Ttbk2^sca11/sca11^* embryos exhibit morphological phenotypes that are strikingly similar to those that we previously described in embryos homozygous for an ENU-induced null allele of *Ttbk2*, *Ttbk2^bby/bby^* [4] (referred to from this point as *Ttbk2^null/null^*), including holoprosencephaly, a pointed midbrain flexure, and randomized heart laterality (**Fig 1A**). The SHH pathway is essential for the patterning of many tissues within the developing embryo, including the neural tube, where a gradient of SHH from the notochord specifies and patterns ventral neural progenitors, including floorplate, V3 interneuron progenitors, and motor neuron progenitors [16]. Similar to *Ttbk2^null/null^*, embryos, the *Ttbk2^sca11/sca11^* embryos exhibited neural patterning defects consistent with a failure to respond to SHH, including the absence of the NKX2.2+ V3 interneuron progenitors that require high levels of SHH activity and ISL1+ motor neurons that are shifted ventrally to span the midline (**Fig 1B vs. E; C vs. F**). We have also examined the phenotype of *Ttbk2^sca11/null^*, confirming that these embryos have the same phenotype (**Fig S1**).

**Figure 1.**
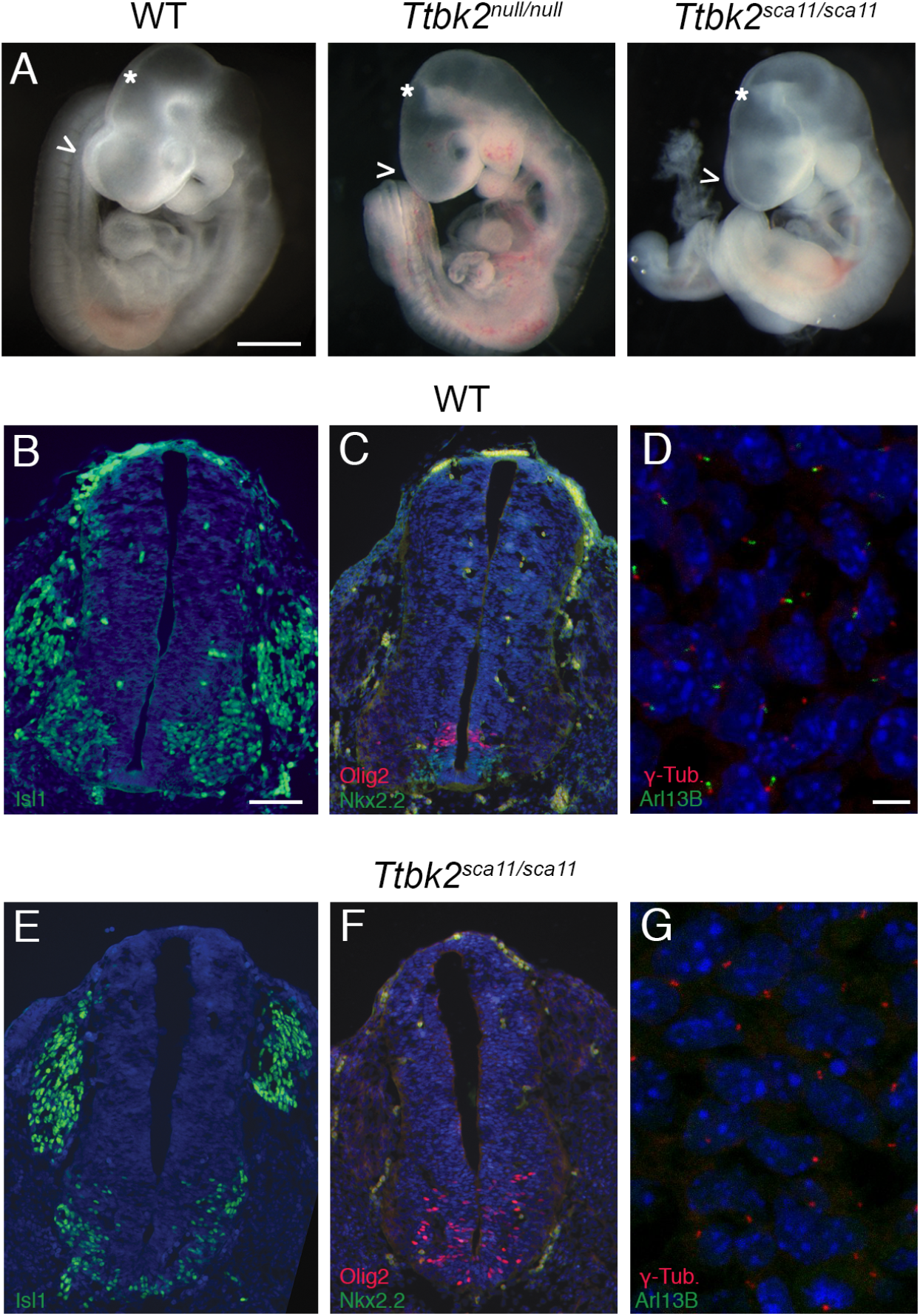
The phenotype of *Ttbk2^sca11/sca11^* embryos is identical to that of the null allele. **(A)** Representative wild-type, *Ttbk2^null/null^*, and *Ttbk2^sca11/sca11^* E10.5 embryos, as indicated. Scale bar = 1mm. Arrowheads point out the forebrain, which in *Ttbk2^null/nul^* and *Ttbk2^sca11/sca11^* fails to form two distinct hemispheres (holoprosencephaly). * Indicates the midbrain flexure which in both mutants is narrowed and takes on a pointed appearance, similar to other mutants with disrupted cilia and SHH signaling. **(B-C, E-F)** Transverse sections through neural tubes of wild-type (B-C) and *Ttbk2^sca11/sca11^* (E-F) E10.5 embryos. Sections were taken at the level of the forelimbs and immunostained for ISL1 (B, E) to label differentiated motor neurons or NKX2.2 and OLIG2 (C, F) to label V3 interneuron progenitors and motor neuron progenitors, respectively. Scale bar = 100μm. **(D, G)** Mesenchymal cells surrounding neural tube of E10.5 WT and *Ttbk2^sca11/sca11^* embryos, immunostained for ARL13B to label cilia (green) and γ-Tubulin (red). Scale bar = 13μm.

*Ttbk2^sca11/sca11^* embryos lacked cilia in mesenchymal cells surrounding the neural tube (**Fig 1D vs. G**), as assayed by immunostaining for the ciliary membrane protein ARL13B. Mouse embryo fibroblasts (MEFs) derived from *Ttbk2^sca11/sca11^* embryos failed to recruit IFT proteins to the basal body and retained the cilium-suppressing centrosomal protein CP110 at the distal mother centriole (**Fig S2**), cellular defects identical to those originally reported for the *Ttbk2^null/null^* allele [4]. Truncated TTBK2 protein produced by the SCA11-associated mutations was reportedly detected in tissues from heterozygous knockin animals [14], however our data indicate that these truncations are unable to function in cilia formation, despite the inclusion of the kinase domain. To better understand how the SCA11-associated mutations might lead to a dominant neurological condition in humans, we next undertook a series of studies to investigate the effect of these truncations on ciliogenesis in the presence of varied amounts of full length TTBK2.

### Decreased rescue of *Ttbk2^sca11/sca11^* MEFs by TTBK2-GFP

Neither *Ttbk2^null/null^* [4], *Ttbk2^sca11/null^*, nor *Ttbk2^sca11/sca11^* embryos form any cilia (**Figs 1**, **S1**, **S2**). However, in cells derived from *Ttbk2^null/null^* embryos, cilia can be fully rescued by expression of WT TTBK2-GFP via retroviral transduction. We previously found that overexpression of TTBK2^SCA11^ in WT fibroblasts using the same method modestly suppresses cilia formation[4]. This led us to propose that *Ttbk2^sca11^* may be a dominant-negative (antimorphic) allele of *Ttbk2*. To test this hypothesis, we expressed WT TTBK2-GFP in MEFs derived from both *Ttbk2^null/null^* and *Ttbk2^sca11/sca11^* embryos using the same retroviral transduction system we previously employed for rescue experiments, and compared the ability of WT TTBK2 to rescue cilia formation in cells of these two genotypes. The frequency of cilia rescue was approximately 2-fold lower in *Ttbk2^sca11/sca11^* MEFs compared to *Ttbk2^null/null^* MEFs (34.1 +/-4.6% vs 66.2 +/-3.3%; p= 0.0002; **Fig 2A, B**). The intensity of ARL13B was also reduced in cilia of *Ttbk2^sca11/sca11^* MEFs expressing TTBK2-GFP compared to cilia in *Ttbk2^null/null^* MEFs (WT: 120.4 +/- 4.96 A.U., *Ttbk2^bby/bby^*+ TTBK2-GFP: 103.2 +/- 3.37 A.U., *Ttbk2^sca11/sca11^*+ TTBK2-GFP: 58.93 +/- 6.14 A.U.; **Fig 2C, D**), although the cilia did not differ significantly in length (WT: 3.468 +/- 0.154, *Ttbk2^bby/bby^*+ TTBK2-GFP: 2.874 +/- 0.074, *Ttbk2^sca11/sca11^*+ TTBK2-GFP: 3.438 +/- 0.194, **Fig 2C, E**). Together, these results suggest that the ability of exogenous TTBK2-GFP to restore cilia in mutant fibroblasts is inhibited by the presence of the truncated TTBK2^SCA11^ protein in these cells.

**Figure 2.**
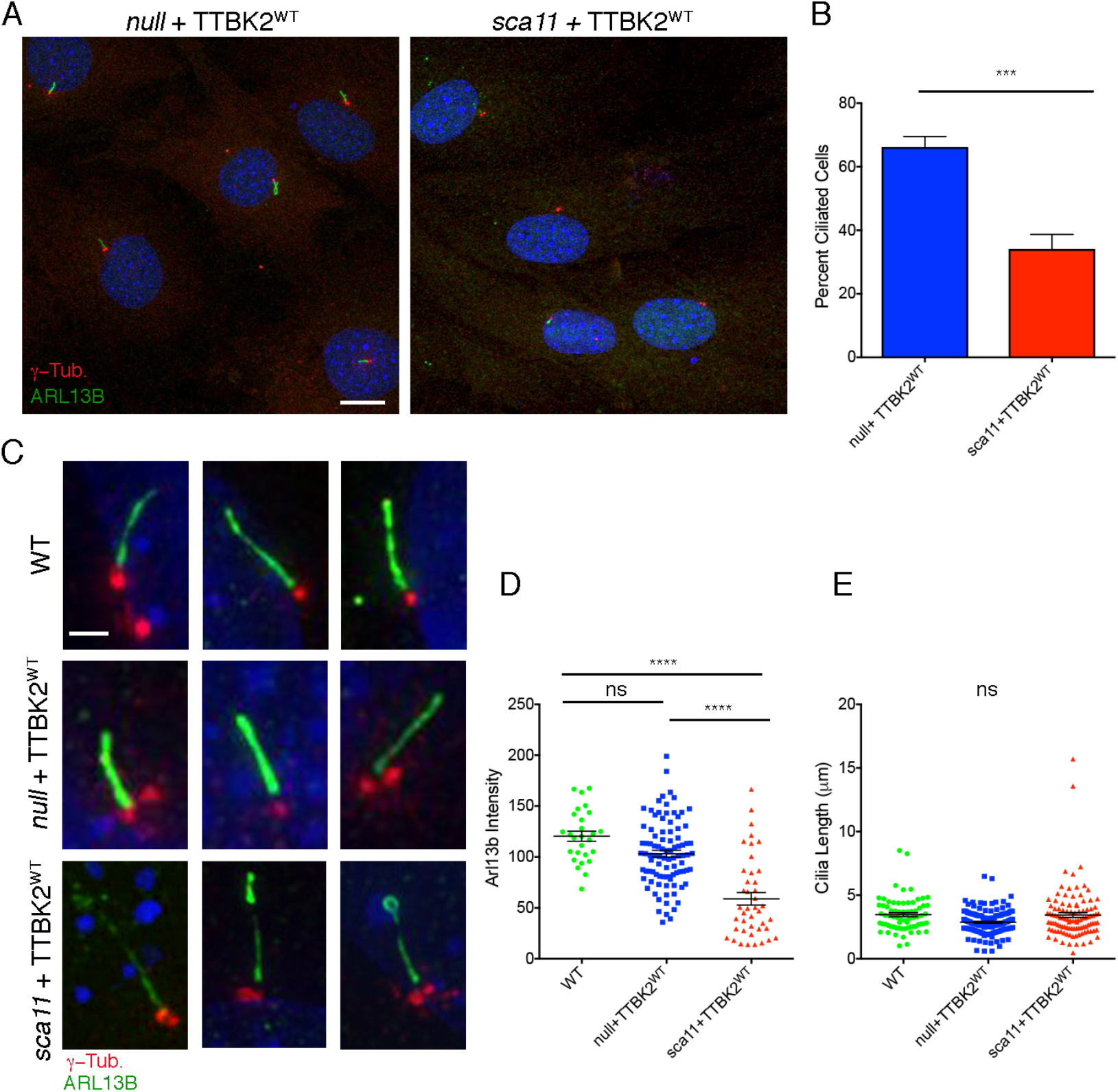
A SCA11-associated allele of *Ttbk2* interferes wild-type TTBK2 function in cilia formation. **(A, B)** A comparison of rescue of cilia formation by WT TTBK2-GFP in MEFs derived from *Ttbk2^null/null^* embryos vs *Ttbk2^sca11/sca11^* embryos. A representative field of cells from each condition is shown in (A) immunostained for ARL13B (green) and γ-Tubulin (red). Scale bar = 20μm. The mean percent of ciliated cells in *Ttbk2^null/null^* MEFs versus *Ttbk2^sca11/sca11^* MEFs after rescue with TTBK2^WT^ is shown in (B). The graph represents the average of 6 fields of cells across two independent experiments (null+TTBK2^WT^ n= 172; sca11+Ttbk2^WT^ n=190). Error bars represent SEM. p=0.0002 by Student T-test. **(C-E)** A comparison of cilia morphology in WT cells, *Ttbk2^null/null^* MEFs rescued with TTBK2-GFP, and *Ttbk2^sca11/sca11^* MEFs rescued with TTBK2-GFP. (C) Depicts two representative images of cilia of each condition immunostained for ARL13B (green) and γ-Tubulin (red). Scale bar = 2μm. ARL13B fluorescence intensity is shown in (D). ARL13b intensity is lower in *Ttbk2^sca11/sca11^*+TTBK2-GFP compared to both WT and *Ttbk2^null/null^* +TTBK2-GFP (1-way ANOVA, p<0.0001), however the mean cilia length is not different between these conditions (E).

### TTBK2^SCA11^ does not physically interact with full length TTBK2

To begin to investigate how the truncated SCA11-associated protein might interfere with the function of WT TTBK2, we tested whether TTBK2^SCA11^ could physically interact with full-length TTBK2. Previous studies have found that TTBK2 molecules physically associate and that TTBK2 can phosphorylate its own C terminus[8], suggesting that like other members of the CK1 family[17], TTBK2 may form a homodimer, and this association could have regulatory significance. We therefore tested the ability of different fragments of TTBK2 (**Fig 3A**) to interact with full-length TTBK2 by co-immunoprecipitation (in all cases GFP or V5 tags were placed at the N terminus of the protein or protein fragment). Consistent with previous reports, V5-tagged full-length TTBK2 co-precipitates with full-length TTBK2-GFP when both constructs are expressed in HEK293T cells (**Fig 3B**). Full-length TTBK2-V5 also co-precipitates with the C-terminus of TTBK2 (TTBK2^306-1243^-GFP), but not with a SCA11-associated TTBK2 truncation (TTBK2^1-443^) (**Fig 3C**). Thus, the C-terminus of TTBK2 (amino acids 450-1243) is essential for this self-interaction. Consistent with this finding, TTBK2^SCA11^-V5 is also unable to co-immunoprecipitate with TTBK2^SCA11^-GFP. The loss of these interactions has implications for the regulation of TTBK2^SCA11^ as well as its interactions with substrates. As the SCA11 truncation does not bind the full-length protein, it is likely that the SCA11 protein acts as a dominant negative by competing with the full-length protein for binding to another protein or proteins.

**Figure 3.**
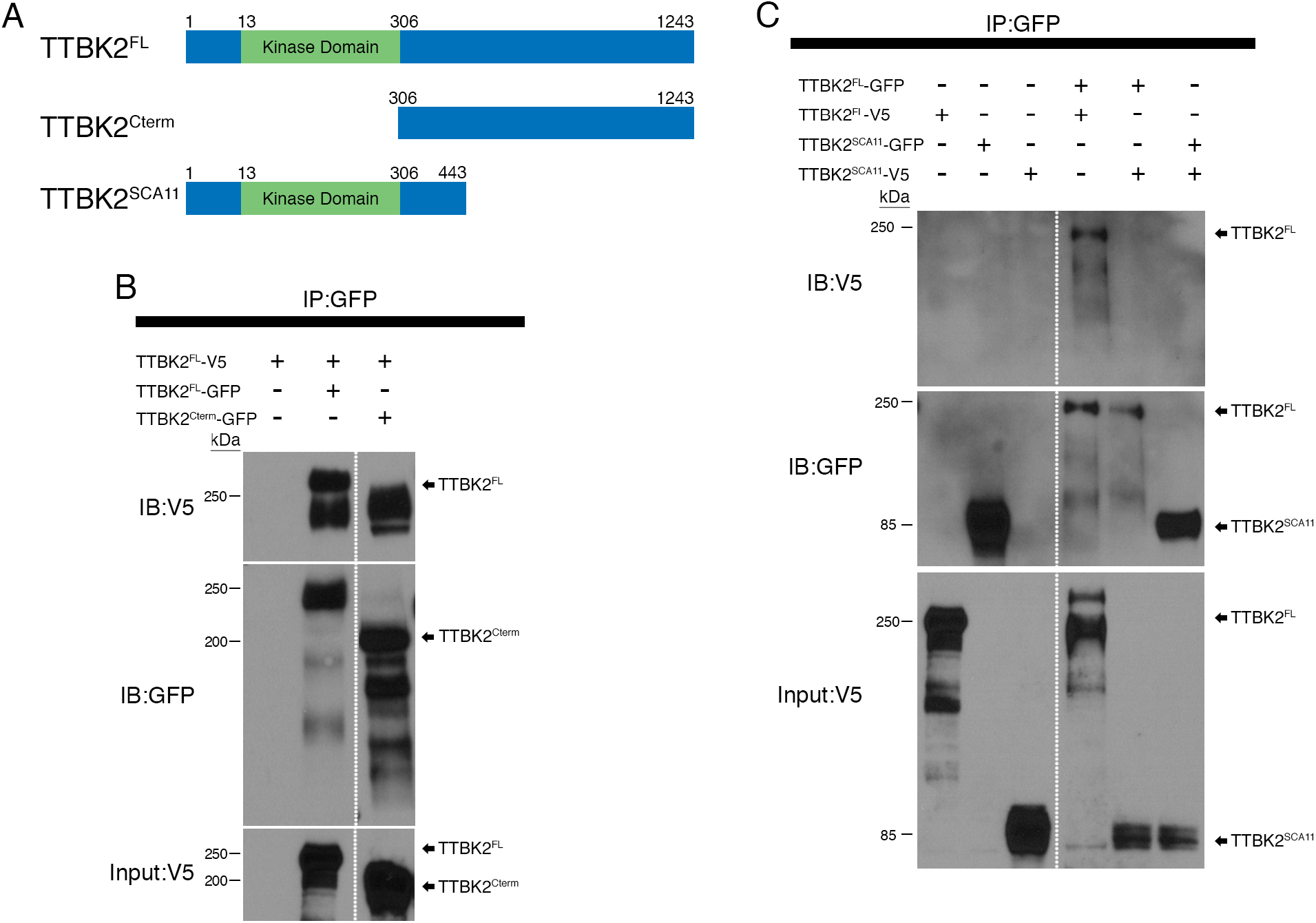
TTBK2 homodimerizes through its C-terminus. **(A)** A schematic representing the various TTBK2 truncations used for co-immunoprecipitation. TTBK2^FL^ corresponds to full length mouse TTBK2. TTBK2^CTerm^ is the amino acids C-terminal to the kinase domain: 306-1243. TTBK2^SCA11^ corresponds to one of the disease-associated mutations, and includes amino acids 1-443. Each is tagged with either V5 or GFP at the N-terminus as indicated in B and C. **(B)** Construct were expressed together as indicated in HEK 293T cells, which were lysed and subjected to immunoprecipitation using anti-GFP conjugated beads. Full length TTBK2 tagged with V5 (TTBK2^FL^-V5) was co-transfected with GFP-tagged full-length TTBK2 (TTBK2^FL^-GFP) or TTBK2^Cterm^. TTBK2^FL^-V5 interacted with TTBK2^FL^-GFP and TTBK2^Cterm^-GFP. Dashed line indicates that the blot image was cropped to remove bands that were not part of the current analysis. Input protein amount was 10% of total lysate for all conditions. **(C)** Constructs encoding either TTBK2^FL^ or TTBK2^SCA11^ tagged with either GFP or V5 as indicated were expressed in HEK293T cells and subjected to immunoprecipitation using anti-GFP conjugated beads. Again, TTBK2^FL^-V5 and TTBK2^FL^-GFP co-precipitated, however TTBK2^FL^ did not co-precipitate with TTBK2^SCA11^-GFP, and TTBK2^SCA11^-V5 did not co-precipitate with TTBK2^SCA11^-GFP. Input protein amount was 10% of total lysate for all conditions. Dashed line indicates that the blot image was cropped to remove bands that were not part of the current analysis. Input protein amount was 10% of total lysate for all conditions.

### *Ttbk2^sca11^* acts as an antimorphic allele

To test genetically whether the SCA11-associated mutations to *Ttbk2* are antimorphic alleles, we again turned to the *Ttbk2^sca11^* knockin mice. Since the human disease SCA11 is an adult-onset phenotype seen in individuals heterozygous for *TTBK2* mutations, we examined the phenotype of adult *Ttbk2^scall/+^* mice. At 3 months of age, the cerebellar architecture of the *Ttbk2^scall/+^* animals was indistinguishable from that of wild type (**Fig S3**). SCA11 in humans reportedly results in mild ataxia, with patients having a normal lifespan and predominantly remaining ambulatory even years after the onset of symptoms[11, 18–20]. We cannot yet rule out that the mice may have subtle or late-onset phenotypes. Another possibility is that the relative dosage of WT or full length TTBK2 required to maintain sufficient ciliary function and/or neuronal function and survival is lower in mice than it is in humans.

To further test whether TTBK2^SCA11^ dominantly interferes with WT TTBK2 function *in vivo*, we next generated an allelic series of *Ttbk2*. We generated mice that carry a gene trap allele of *Ttbk2* (*Ttbk2^tm1a(EUCOMM)Hmgu^*) from ES cells obtained from the European Mutant Mouse Cell Repository (EuMMCR). Although the targeting strategy was designed to trap splicing of an early *Ttbk2* exon (**Fig S4A**), the homozygous gene trap mice (*Ttbk2^gt/gt^*) were viable past weaning, developing variably penetrant hydrocephalus and polycystic kidneys by 6 months of age (**Fig S4C, D**). Transcript analysis showed that this allele produced mRNAs with the predicted gene trap transcript and a wild-type RNA formed by splicing around the gene trap insertion (**Fig S3B**). Consistent with this, by Western blot we detected a small amount of TTBK2 protein, running at the same molecular weight as WT TTBK2 (**Fig S3E**). We conclude that *Ttbk2^gt^* is a partial loss-of-function (hypomorphic) allele that produces a reduced amount of wild-type, full-length TTBK2 protein.

Consistent with the hypomorphic character of the gene trap allele,*Ttbk2^null/gt^* embryos had a phenotype intermediate between that of *Ttbk2^null/null^* and the *Ttbk2^gt/gt^* homozygotes: *Ttbk2^null/gt^* embryos and neonates were recovered at nearly Mendelian frequencies up to birth (P0) but died by P1 (**Table S1**). At E15.5, in contrast to *Ttbk2^gt/gt^* embryos, which showed wild-type morphology, *Ttbk2^null/gt^* embryos had fully penetrant polydactyly on all 4 limbs, consistent with a disruption in Hh-dependent limb patterning **(Fig 4A-C, Table S2**).

**Figure 4.**
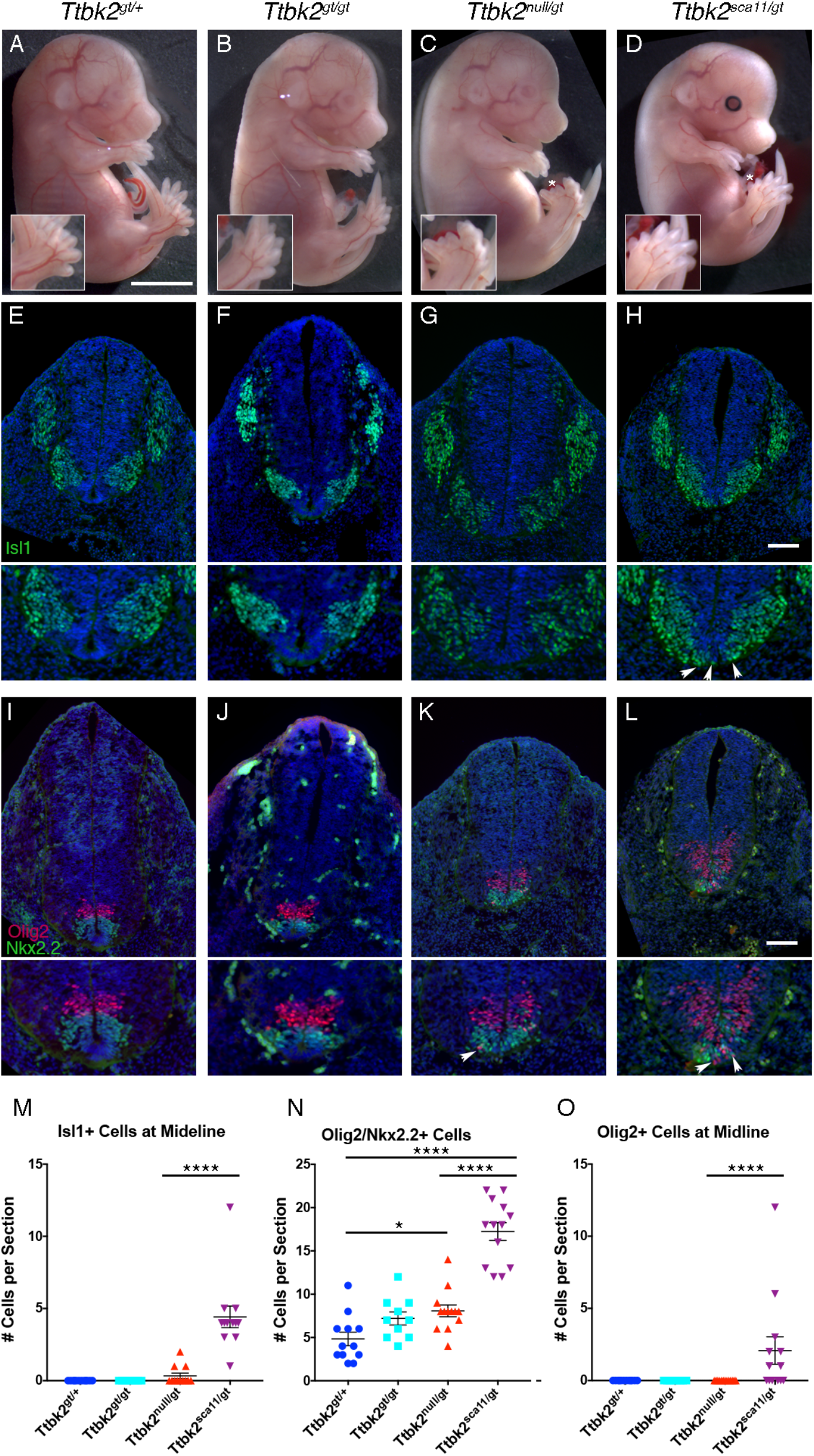
A *Ttbk2* allelic series highlights reveals exacerbated SHH-related phenotypes in *Ttbk2^sca11/gt^* embryos. **(A-D)** Representative E15.5 embryos of each of the indicated *Ttbk2* allelic combinations. Scale bar = 3mm. * Denotes duplicated digits. Insets are 1.5X enlargements of the hindlimbs shown in each image. **(E-H)** Transverse sections through the neural tube of E10.5 embryos of the indicated genotypes. Sections were taken at the forelimb level and immunostained for the motor neuron marker ISL1. Scale bar = 100μm. Images below are 1.5X enlargements of the ventral portion of each neural tube section. Arrowheads in H denote ventrally displaced motor neurons spanning the ventral midline. **(I-L)** Representative transverse sections through E10.5 neural tubes of embryos of each indicated genotype immunostained for NKX2.2 (green) to label V3 interneuron progenitors and OLIG2 (red) to label motor neuron progenitors. Scale bar = 100μm. Images below are 1.5X enlargements of the ventral portion of each neural tube section. Arrows in K and L denote ventrally displaced OLIG2+ progenitors. **(M)** Quantification of the number of ISL1+ cells seen at the ventral midline per section. Sections were taken at the forelimb level as pictured in E-H. 3 sections in 4 different embryos were for each genotype except for *Ttbk2^gt/gt^* in which 3 embryos were evaluated. Each point represents the count for one section. ISL1+ cells were never observed at the midline for *Ttbk2^gt/+^* or *Ttbk2^gt/gt^*. By contrast, in *Ttbk2^null/gt^*, 3 out of 4 embryos evaluated had a small number of cells at the midline in at least 1 section, and in *Ttbk2^sca11/gt^* embryos, ISL1+ cells were counted at the midline in all sections of all embryos. The number of ISL1+ is significantly higher in *Ttbk2^sca11/gt^* embryos than *Ttbk2^null/gt^* (1-way ANOVA, p<0.0001). Error bars denote SEM. **(N)** Quantification of cells double positive for both OLIG2 and NKX2.2 per section. Sections were taken at the forelimb level as pictured in I-L, and numbers evaluated were as in M. Significantly more double positive cells were observed in *Ttbk2^null/gt^* and *Ttbk2^sca11/gt^* relative to *Ttbk2^gt/+^* (1-way ANOVA, p=0.036 and p<0.0001, respectively). In addition more double positive cells were counted in *Ttbk2^sca11/gt^* embryos compared with *Ttbk2^null/gt^* (p< 0.0001). Error bars denote SEM. **(O)** Quantification of the number of OLIG2+ cells counted at the ventral midline per section. Sections were taken at the forelimb level as pictured in I-L, and numbers evaluated were as in M. OLIG2+ cells were only observed in *Ttbk2^sca11/gt^*, and 4/4 embryos evaluated had OLIG2+ midline cells in at least one section. This was a statistically significant increase relative to each of the other genotypes (1-way ANOVA, p< 0.0001). Error bars denote SEM.

We reasoned that the *Ttbk2^gt^* allele, with lowered levels of TTBK2 protein, might provide a sensitized genetic background to better compare the effects of the *Ttbk2^null^* and *Ttbk2^sca11^* alleles. *Ttbk2^sca11/gt^* embryos showed similar overall morphology to *Ttbk2^null/gt^* embryos at E15.5, with fully penetrant polydactyly on all 4 limbs. While some *Ttbk2^sca11/gt^* neonates were recovered at P0, they were present at a sub-Mendelian frequency: only 9.5% of pups (6/63 compared with 16/63 expected; p= 0.0189) recovered at birth from *Ttbk2^gt/+^* x *Ttbk2^sca11/+^* crosses genotyped as *Ttbk2^sca11/gt^* (summarized in **Tables S1, S2**), suggesting some prenatal lethality.

We compared ventral neural patterning in *Ttbk2^gt/gt^*, *Ttbk2^null/gt^*, and *Ttbk2^sca11/gt^* embryos to determine whether *Ttbk2^sca11/gt^* embryos had more severe disruption of SHH signaling than seen in *Ttbk2^null/gt^* embryos. We found that neural patterning in E10.5 *Ttbk2^gt/gt^* embryos was similar to that in *Ttbk2^gt/+^* embryos (**Fig 4 E-F, I-J**), and *Ttbk2^null/gt^* embryos exhibited only mild defects in neural patterning. In these mutants, the distribution of motor neurons, labeled with Islet1 (ISL1) was very similar to that of *Ttbk2^gt/+^* though a small number of motor neurons were found at the ventral midline (**Fig 4G, M**). We also observed an increase in the number of cells positive for both NKX2.2 and OLIG2 relative to *Ttbk2^gt/+^*(**Fig 4K, N**). In contrast, the ISL1+ motor neuron domain was shifted ventrally in *Ttbk2^sca11/gt^* embryos and ISL1+ cells were found at the ventral midline in all sections examined (**Fig 4H, M**). We also observed extensive intermixing of OLIG2+ and NKX2.2+ progenitor populations, with a larger number of cells positive for both NKX2.2 and OLIG2 compared to other genotypes (**Fig 4L, N**). In addition, OLIG2+ cells were often found at the ventral midline in the *Ttbk2^sca11/gt^* embryos, in contrast with the other genotypes in which this dramatic ventral shift was not observed (**Fig 4L, O**). Together, these data are consistent with a more severe disruption in SHH-dependent patterning in the *Ttbk2^sca11/gt^* embryos. This enhanced SHH patterning phenotype, combined with the increase in embryonic lethality of the *Ttbk2^sca11/gt^* animals, provides genetic support for *Ttbk2^sca11^* as a dominant negative allele.

### TTBK2 controls cilia length, trafficking and stability

To assess whether the more severe developmental defects in *Ttbk2^sca11/gt^* embryos were due to greater defects in ciliary trafficking, structure, or stability, we analyzed cilia in MEFs derived from embryos of each genotype of the *Ttbk2* allelic series. Following serum starvation, we found that a mean of 69.1 +/- 3.64 % of *Ttbk2^gt/+^* cells were ciliated (**Fig 5A, J**), whereas in *Ttbk2^gt/gt^* and *Ttbk2^null/gt^* an average of 45.9 +/- 3.66 % and 43.8 +/- 3.35 % of cells were ciliated, respectively (*Ttbk2^gt/+^* vs *Ttbk2^gt/gt^* p =0.0003; *Ttbk2^gt/+^* vs *Ttbk2^null/gt^* p<0.0001; *Ttbk2^gt/gt^* vs *Ttbk2^null/gt^* p= 0.9772; **Fig 5B-C, J**). There were clearly fewer cilia in *Ttbk2^sca11/gt^* cells, with an average of 18.9 +/- 3.65% of cells having a cilium (*Ttbk2^null/gt^* vs *Ttbk2^sca11/gt^* p<0.0001; **Fig 5D, J**). These findings suggest that the increased severity of the embryonic phenotypes correlates with a decrease in cilia number. While the mean cilia length was reduced in all of the *Ttbk2* mutants relative to *Ttbk2^gt/+^* cells, cilia length did not differ significantly between the different mutant allelic combinations (**Fig 5K**). We observe similar overall trends examining the cilia of the embryo (mesenchymal tissue surrounding the neural tube) where the frequency of cilia is reduced in the hypomorphic mutants, and more dramatically reduced in *Ttbk2^sca11/gt^* (**Fig S5**).

**Figure 5.**
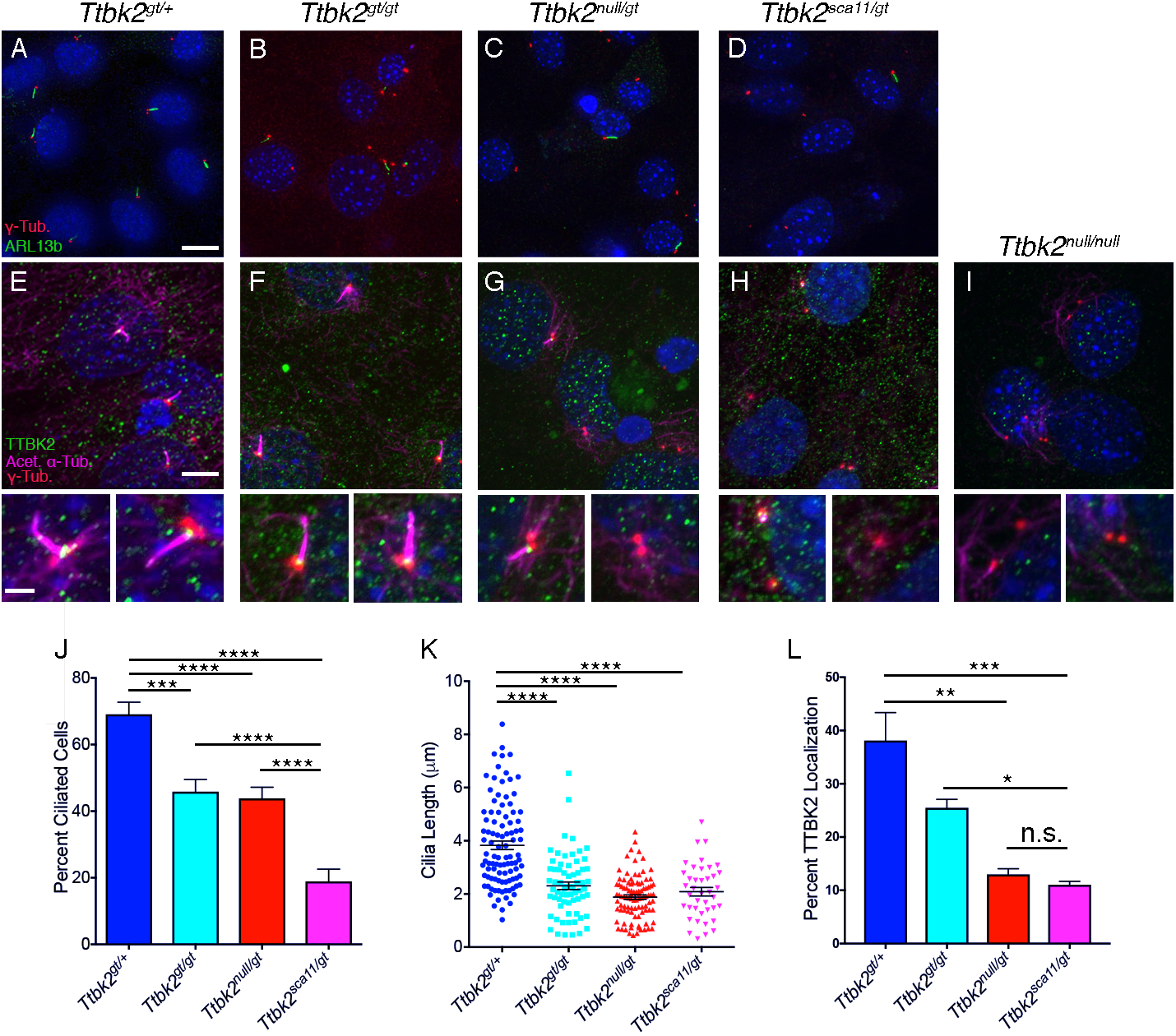
Ciliary defects are evident in *Ttbk2* hypomorphic mutant cells. **(A-D)** Representative images of MEFs taken from the embryos of the indicated and serum starved for 48-hours to induce ciliogenesis. Cilia were immunostained for ARL13b (green) to label cilia and γ-Tubulin (red) to label centrosomes. Scale bar = 20μm. **(E-I)** Localization of endogenous TTBK2 (green) in MEFs of each indicated genotype. Cells were counterstained with γ-Tubulin (red) to label centrosomes and Acetylated α-Tubulin (magenta) to label cilia. Staining of *Ttbk2^null/null^* cells is shown as a negative control. Lower panels show zoomed images of from individual cells seen in the top panel. Scale bar = 10μm, 2μm for zoomed panels. **(J)** Quantification of the percentage of ciliated cells from MEFs of each genotype following 48 hours of serum starvation. Cilia frequency is reduced in all mutants compared with heterozygous cells, and is further reduced in *Ttbk2^sca11/gt^* cells relative to *Ttbk2^null/gt^*. Bars represent the mean percentage of ciliated cells across 3 experiments with 3 replicates each (n= 289, *Ttbk2^gt/+^*; n= 252, *Ttbk2^gt/gt^*; n= 584, *Ttbk2^null/gt^*; n= 340 *Ttbk2^sca11/gt^*). Error bars denote SEM. **(K)** Quantification of cilia length in MEFS of each genotype following 48 hours of serum starvation. Cilia in each of the mutant MEFs were significantly shorter than heterozygous cells, but did not differ significantly from each other. **(L)** Quantification of the percentage of cells with endogenous TTBK2 localized to the centrosome following 48 hours of serum withdrawal. *Ttbk2* mutant cells, particularly *Ttbk2^null/gt^* and *Ttbk2^sca11/gt^* have reduced TTBK2 at the centrosome, however these two genotypes did not differ from one another with respect to TTBK2 localization. For J-L, stars denote statistical comparisons between the indicated groups (1- way ANOVA with Tukey-Kramer post hoc test, **** denotes p< 0.0001, *** denotes p<0.001, ** denotes p=0.0017, * denotes p= 0.014.)

We also examined the percentage of cells with endogenous TTBK2 at the mother centriole or basal body in MEFs derived from each genotype. Relative to*Ttbk2^gt/+^,* the percentage of cells with TTBK2 localized at the mother centriole/basal body in *Ttbk2^gt/gt^* trended towards being slightly reduced, though this was not statistically significant (**Fig 5E--F, L**; 38.1 +/- 9.11% for *Ttbk2^gt/+^* cells vs 25.5 +/- 2.76% for *Ttbk2^gt/gt^*; p= 0.052). As expected, centriolar TTBK2 was further reduced in *Ttbk2^null/gt^* and *Ttbk2^sca11/gt^* cells (**Fig 5G-H, L**; 13.0 +/- 1.86% and 11.1 +/- 1.12%, respectively; *Ttbk2^gt/+^* vs *Ttbk2^null/gt^* p=0.001; *Ttbk2^gt/+^* vs *Ttbk2^sca11/gt^* p= 0.0006), but there was no significant difference between these two genotypes (**Fig 5L**; p= 0.9608), implying that the presence of TTBK2^SCA11^ does not interfere with full length TTBK2 function by impairing its localization to the presumptive basal body.

Since our data indicate that hypomorphic *Ttbk2* mutants have shorter cilia in addition to forming cilia at a reduced frequency, we hypothesized that TTBK2 may be required for ciliary trafficking and stability as well as for the initiation of ciliogenesis. To further investigate the role of TTBK2 in cilia structure and/or trafficking following initial assembly of the axoneme, we examined trafficking of HH pathway components in the cilia of MEFS of each genotype. The transmembrane protein SMO is critical for HH pathway activation, and becomes enriched within the cilium upon stimulation of the pathway with SHH or various agonists [21,22]. We found that the amount of SMO in the cilium upon stimulation of cells with SMO agonist (SAG) was comparable between *Ttbk2^gt/+^* cells and either *Ttbk2^gt/gt^* or *Ttbk2^null/gt^* cells, as measured by average intensity of SMO within the Acetylated α-Tubulin+ cilium (mean intensity of 82.2 +/- 3.43, 89.2 +/- 5.3, and 78.9 +/- 4.65 A.U., respectively). In contrast, SMO intensity was clearly reduced in the axonemes of *Ttbk2^sca11/gt^* cells (mean intensity of 51.0 +/- 3.78 A.U.) relative to *Ttbk2^null/gt^* (p= 0.0003), as well as each of the other genotypes (vs *Ttbk2^gt/+^* and *Ttbk2^gt/gt^* p<0.0001), consistent with the exacerbated SHH signaling-related phenotypes observed in these mutant embryos (**Fig 6A, B**).

We also examined the trafficking of other HH pathway components within cilia in response to pathway activation. GLI2 is a transcription factor that mediates activation of target genes in response to HH ligands. GLI2 localizes to the tips of cilia, and becomes strongly enriched at the cilium tip in response to HH pathway activation[23] by SHH or SAG. There was no difference in GLI2 ciliary tip localization or intensity in response to SAG between *Ttbk2^gt/+^* and any of the mutant alleles (**Fig 6C, D**). KIF7 is the vertebrate homolog of the *Drosophila* protein COS2 and essential for the establishment and maintenance of the microtubule structure of the cilium in mammals, and for the stability of the axoneme[24,25]. Like GLI2, KIF7 normally becomes enriched at the tips of cilia in response to SAG, however in contrast to the results with GLI2, the percentage of cells with KIF7 localized to the tip of the cilium in the presence of SAG was significantly reduced in *Ttbk2^sca11/gt^* mutants relative to other genotypes (*Ttbk2^gt/+^*: 82.2 +/- 2.16%, *Ttbk2^gt/gt^*: 75.6 +/- 1.73%, *Ttbk2^null/gt^*: 62.6 +/- 7.93%, *Ttbk2^sca11/gt^*: 37.8 +/- 1.1%; **Fig 6E, F,**), and was clearly less than in *Ttbk2^null/gt^* (p= 0.012). Thus, consistent with the more severe SHH-related patterning phenotypes observed in the *Ttbk2^sca11/gt^* embryos, trafficking of a subset of signaling molecules is impaired in cells of this genotype, consistent with possible disruptions in ciliary trafficking.

**Figure 6.**
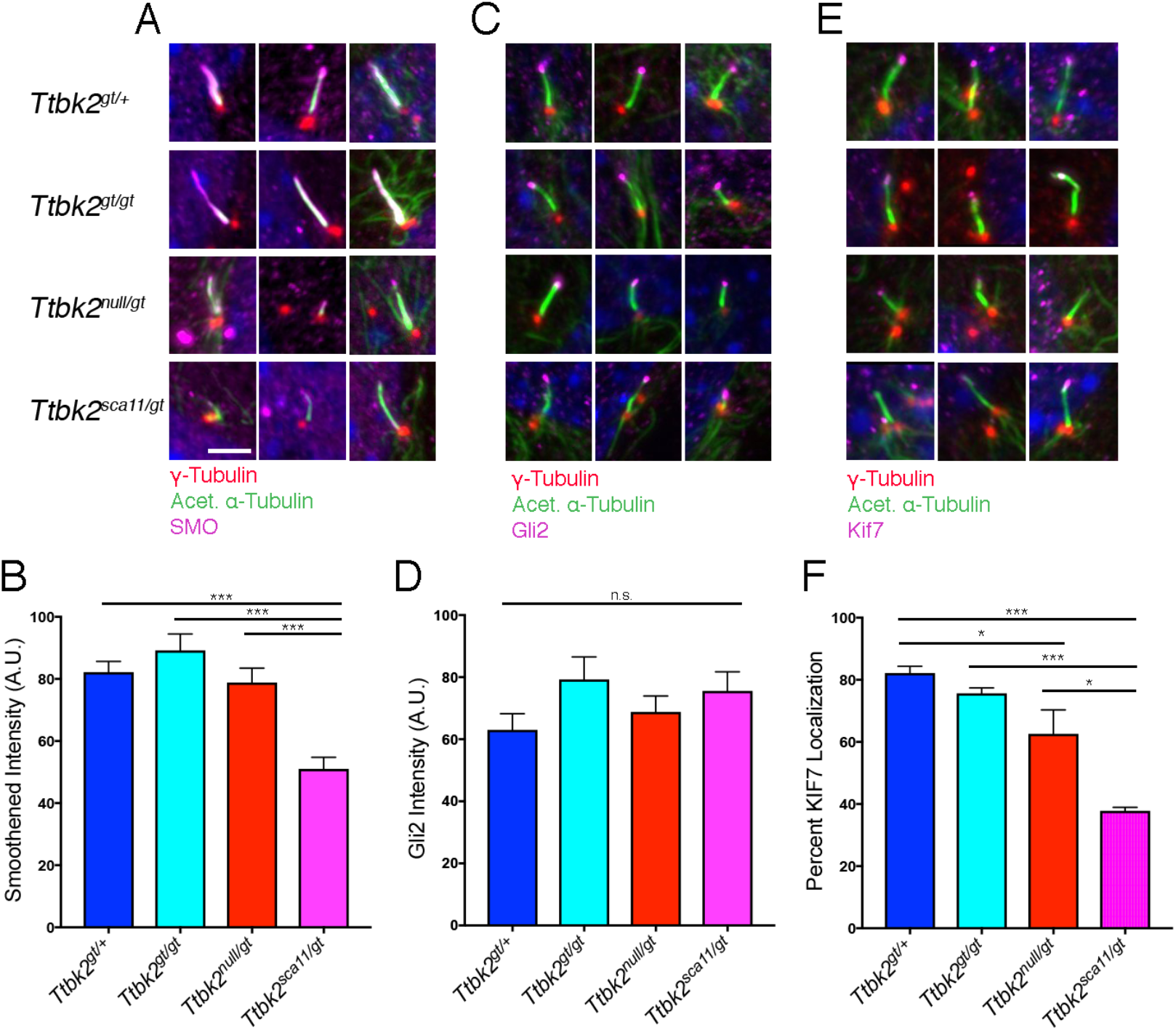
Localization of key ciliary proteins is impaired in *Ttbk2^sca11/gt^* cilia. **(A, B)** SMO localization in heterozygous and *Ttbk2* mutant MEFs. Immunostaining for SMO (magenta) is shown for each of the indicated genotypes in (A), with centrosomes stained for γ- Tubulin (red), and the ciliary axoneme labeled with Acetylated α-Tubulin (green). Cells were serum starved for 24 hours before being treated with 200nm SAG for an additional 24 hours. Two representative images are shown. (B) depicts a quantification of pixel intensity of SMO within the ciliary axoneme of each genotype upon stimulation with SAG. Bars represent the mean intensity of 50 measurements across 3 replicates for each genotype. Error bars depict SEM. Statistical comparison was performed by 1-way ANOVA with Tukey-Kramer post-hoc test. *Ttbk2^sca11/gt^* cells display reduced SMO intensity within the cilium compared with each of the other genotypes (p<0.0001 vs *Ttbk2^gt/+^* and *Ttbk2^gt/gt^*; p= 0.0003 vs *Ttbk2^null/gt^*). **(C,D)** Localization of GLI2 (magenta) to the ciliary tip in MEFs of each indicated genotype. In (C) cells are counterstained for γ-Tubulin (red) to label centrosomes and Acetylated α-Tubulin (green) to label the axonemes. Three representative images are shown for each genotype. Quantification of pixel intensity of GLI2 at the ciliary tip of each genotype is shown in (D). **(E,F)** Localization of KIF7 (magenta) to the ciliary tip in MEFs of each indicated genotype. In (E) cells are counterstained for γ-Tubulin (red) to label centrosomes and Acetylated α-Tubulin (green) to label the axonemes. Three representative images are shown for each genotype. The percentage of cilia with KIF7 localized to the ciliary tip upon treatment with SAG is shown in (F). We find that the frequency of KIF7 localization is reduced in *Ttbk2^sca11/gt^* cells compared with other genotypes (1-way ANOVA, p= 0.0003 vs *Ttbk2^gt/+^*; p=0.0009 vs *Ttbk2^gt/gt^*; p=0.012 vs *Ttbk2null/gt*.) n=50 cilia pooled from 3 biological replicates. Scale bar = 5μm

To further investigate whether ciliary trafficking is disrupted *Ttbk2* hypomorphic mutant cells, and whether this is exacerbated in *Ttbk2^sca11/gt^* cells in particular, we assessed other factors that control cilia trafficking and stability[25]. Since the shorter cilia observed for each of the *Ttbk2* hypomorphic allele combinations relative to *Ttbk2^gt/+^* cells could be due to defects in the protein machinery that mediates assembly of the ciliary axoneme, the intraflagellar transport (IFT) machinery, we examined the localization of IFT components in MEFs of each genotype. We measured the average intensity of IFT81, IFT88, and IFT140 within the ciliary axoneme of MEFs of each genotype. For IFT81, the average intensity was not significantly changed within the axoneme in *Ttbk2^bby/gt^* and *Ttbk2^sca11/gt^* cells relative to the other genotypes (**Fig 7A, B;** *Ttbk2^gt/+^*: 11.49 +/- 0.68 AU, *Ttbk2^gt/gt^*: 13.15 +/- 0.88 AU, *Ttbk2^null/gt^*: 16.07 +/- 1.93 AU, *Ttbk2^sca11/gt^*: 15.49 +/- 1.56 AU). For IFT88, average intensity within the axoneme also varied only modestly by genotype, with modest increases in the average intensity within the axoneme in *Ttbk2^null/gt^* and *Ttbk2^sca11/gt^* relative to the other genotypes (**Fig 7D, E;** *Ttbk2^gt/+^*: 40.54 +/- 1.73 AU, *Ttbk2^gt/gt^*: 45.06 +/- 2.55 AU, *Ttbk2^null/gt^*: 65.61 +/- 3.73 AU, *Ttbk2^sca11/gt^*: 51.89 +/- 3.00 AU). For IFT140 the average intensity was reduced in the axoneme of *Ttbk2^sca11/gt^* cells relative to other genotypes (**Fig 7G, H**; *Ttbk2^gt/+^*: 54.4 +/-3.58 AU, *Ttbk2^gt/gt^*: 65.55 +/- 6.11 AU, *Ttbk2^null/gt^*: 74.62 +/- 7.74 AU, *Ttbk2^sca11/gt^*: 37.13 +/- 3.26 AU). Thus, we identified a specific reduction in IFT-A machinery to the axonemes in *Ttbk2^sca11/gt^* cells relative to other genotypes.

**Figure 7.**
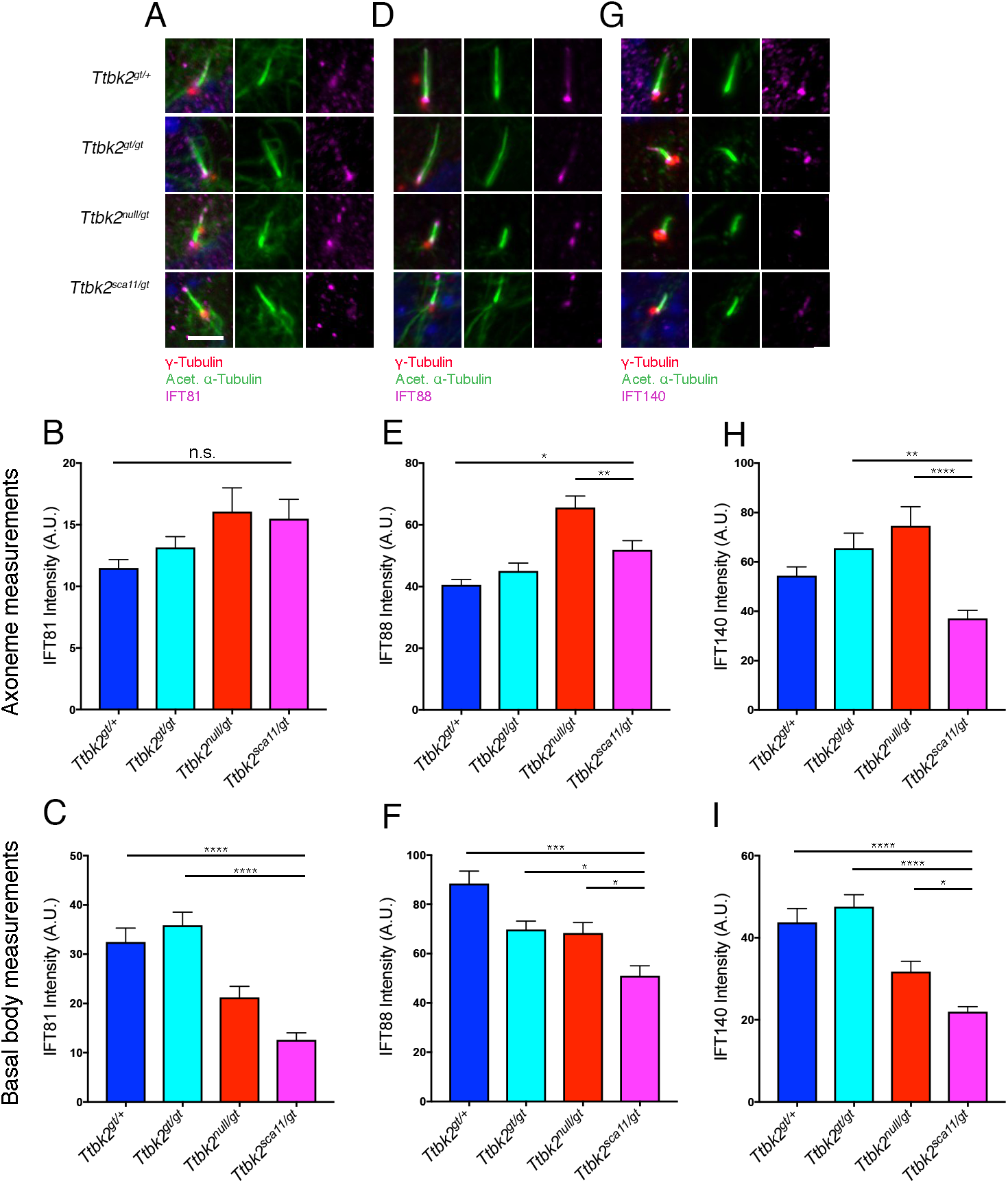
Trafficking defects in the *Ttbk2* allelic series cilia. **(A-C)** Immunostaining in MEFs of each indicated genotype for IFT81 (magenta), γ-Tubulin (red) to label centrosomes and Acetylated α-Tubulin (green) to label the cilia (A). Quantification of IFT81 pixel intensity throughout ciliary axoneme (B) or the basal body pool only (C). Quantification of pixel intensity throughout the axoneme is shown in (B) but was not statistically significant between indicated genotypes. IFT81 pools are depleted in *Ttbk2^sca11/gt^* basal bodies compared to other genotypes (C). Statistical comparison was performed by 1-way ANOVA with Tukey-Kramer post-hoc test. (p<0.0001 vs *Ttbk2^gt/+^*; p=0.0106 vs *Ttbk2^gt/gt^*; p=0.0150 vs *Ttbk2^null/gt^*). Scale bar = 5μm. **(D-F)** Immunostaining of IFT88 (magenta) shows depleted staining throughout the axoneme and at the basal body for *Ttbk2^sca11/gt^* cells (D). Quantification of pixel intensity throughout the axoneme is shown in (E). Statistical comparison was performed by 1-way ANOVA with Tukey-Kramer post-hoc test. (p=0.0282 vs *Ttbk2^gt/+^*; p=0.0052 vs *Ttbk2^null/gt^*).Quantification of pixel intensity at the basal body is shown in (F). Statistical comparison was performed by 1-way ANOVA with Tukey-Kramer post-hoc test. (p<0.0001 vs *Ttbk2^gt/+^*; p<0.0001 vs *Ttbk2^gt/gt^*; p=0.0647 vs Ttbk2^null/gt^). **(G-I)** Immunostaining of IFT140 (magenta) shows depleted staining at the basal body of the axonemes for *Ttbk2^sca11/gt^* cells(G). Quantification of pixel intensity throughout the axoneme is shown in (H). *Ttbk2^sca11/gt^* axonemes display less IFT140 compared to other other genotypes. Statistical comparison was performed by 1-way ANOVA with Tukey-Kramer post-hoc test. (p=0.0022 vs *Ttbk2^gt/gt^*; p<0.0001 vs Ttbk2^null/gt^). Quantification of pixel intensity at the basal body is shown in (I). Statistical comparison was performed by 1-way ANOVA with Tukey-Kramer post-hoc test. (p<0.0001 vs *Ttbk2^gt/+^*; p<0.0001 vs *Ttbk2^gt/gt^*; p=0.0484 vs *Ttbk2^null/gt^*). n=50 or more basal bodies and cilia pooled from 3 biological replicates.

We also separately examined the average intensity of IFT81, IFT88 and IFT140 at the basal body in cells of each genotype. We found that for each of the IFT proteins we examined, this basal body pool of IFT proteins was significantly reduced in average intensity in the *Ttbk2^null/gt^* and *Ttbk2^sca11/gt^* cells relative to *Ttbk2^gt/+^* or *Ttbk2^gt/gt^*. For both IFT88 and IFT140, average intensity at the basal body was significantly lower in *Ttbk2^sca11/gt^* cells than in *Ttbk2^null/gt^* (**Fig 7C, F, I**).

Taken together, this data suggests that reduced function of TTBK2 affects the localization of IFT components, particularly in the pools of IFT that form at the basal body, with the amount and distribution of IFT proteins within the axoneme affected to a lesser degree. These findings are aligned with our prior work showing that the basal body pools of IFT proteins are lost in *Ttbk2^null/null^* cells, a defect that is thus far specific to TTBK2 and to proteins such as CEP164 that act upstream of TTBK2 to mediate its localization to the distal appendages.

The defects we observed in the *Ttbk2* hypomorphic mutants with respect to changes in IFT proteins as well as impaired trafficking of SMO and KIF7 led us to hypothesize that the cilia of these mutants may have defects in their structure and/or stability. Post-translational modifications of axonemal microtubules are often impaired in mutants, such as *Kif7*, in which the structure and stability of the axoneme are disrupted [25]. We have therefore examined both acetylation and glutamylation of tubulin, two modifications associated with stability of the ciliary axoneme [26]. We have not observed any alterations in the acetylation of microtubules within the cilia of the *Ttbk2* hypomorphic mutants, however we observe a marked effect on tubulin polyglutamylation, which is important for establishing ciliary structure and length [27,28]. Intensity of polyglutamylated tubulin within the cilium was comparable between *Ttbk2^gt/+^* and *Ttbk2^gt/gt^* cells (mean intensity of 111.8 +/- 5.42 and 93.64 +/- 5.66 A.U. respectively) but was significantly reduced in *Ttbk2^null/gt^* cells (mean intensity of 75.8 +/- 4.85 A.U.) and further reduced in *Ttbk2^sca11/gt^* cells (mean intensity of 55.1 +/- 4.32 A.U.; **Fig 8A, B**). Reduction of tubulin polyglutamylation is associated with defects in cilium assembly and stability in a variety of organisms [27–29], and recent evidence also suggests that hypo-glutamylation of ciliary microtubules promotes disassembly of primary cilia and impairs the trafficking of signaling molecules within the axoneme [30].

**Figure 8.**
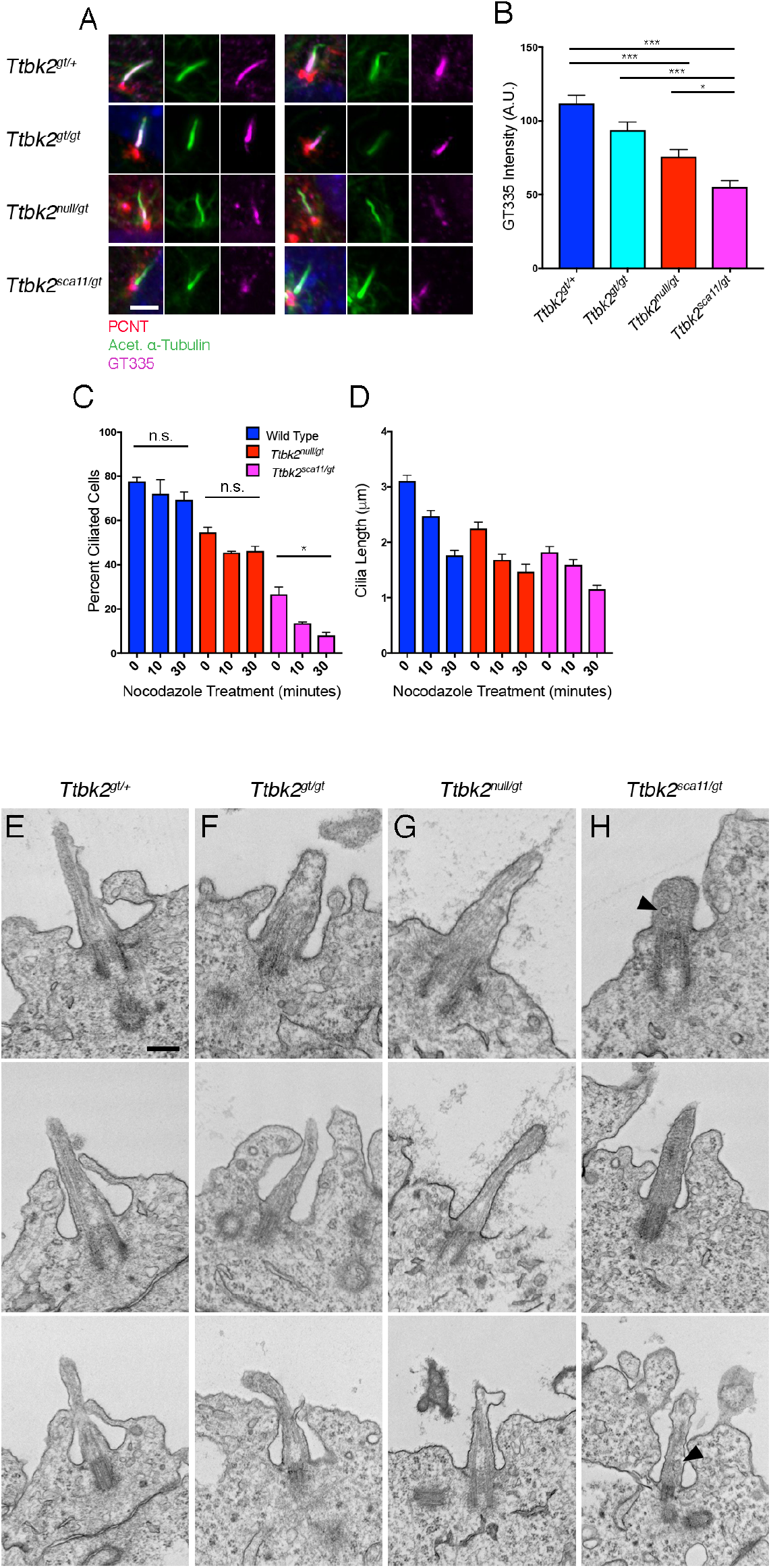
Microtubule dynamics are impaired in *Ttbk2^sca11/gt^* cilia. **(A-B)** Polyglutamylated tubulin localization in cilia of MEFs derived from embryos of the indicated genotypes. Immunostaining for GT335, which recognizes polyglutamylated tubulin (magenta) as well as PCNT (red) to label centrosomes and Acetylated α-Tubulin (green) to label cilia is shown in (A). A quantification of the pixel intensity of GT335 within cilia is shown in (B). Polyglutamylated tubulin levels are reduced in *Ttbk2^sca11/gt^* relative to the other genotypes of the allelic series (p<0.0001 vs vs *Ttbk2^gt/+^*; p<0.0001 vs vs *Ttbk2^gt/gt^*; p= 0.0327 vs *Ttbk2^null/gt^*). Scale bar = 5μm. **(C-D)** Wild type, *Ttbk2^null/gt^*, and *Ttbk2^sca11/gt^* MEFs were treated with 10μM Nocodazole for 10-30 minutes and percentage of ciliated cells was determined. While in WT and *Ttbk2^null/gt^* cells the percentage of ciliated cells reduced only slightly within 30 minutes, cilia were rapidly lost in *Ttbk2^sca11/gt^* MEFs upon treatment with nocodazole (p= 0.0112) (C). Cilia length was determined throughout nocodazole treatment for Wild type, *Ttbk2^null/gt^*, and *Ttbk2^sca11/gt^*. As expected, cilia shortened over time, though not significantly between genotypes over time. n=50 cilia pooled from 3 biological replicates. **(E-H)** Representative Transmission Electron Microscopy (TEM) images of cilia in the embryonic neural tube from e10.5 embryos of the indicated genotypes. Additionally, vesicles can be seen in *Ttbk2^sca11/gt^* axonemes (arrow heads, H) which were not present in other genotypes. Scale bar = 0.5μm

To further examine cilium stability across the *Ttbk2* allelic series, we treated MEFs derived from embryos of each genotype with nocodazole. Because the microtubule doublets of the ciliary axoneme are more stable than cytoplasmic microtubules, treatment of WT cells with nocodazole for a short period has a limited effect on cilia length or frequency [25]. After treatment of MEFs with nocodazole for 10 or 30 minutes, the percentage of ciliated cells in WT or *Ttbk2^null/gt^* cells decreased modestly (**Fig 8C**; for WT, 77.7 +/- 1.73% of cells were ciliated at T0, 72.2 +/- 6.33% at 10 minutes, and 69.3 +/- 3.5% at 30 minutes; for *Ttbk2^null/gt^*, 54.6 +/- 2.31% of cells were ciliated at T0, 45.5 +/- 0.64% at 10 minutes, and 46.3 +/- 1.96% at 30 minutes). In contrast, in *Ttbk2^sca11/gt^* cells treatment with nocodazole caused a rapid reduction in ciliated cells (from 26.6 +/- 3.38% at T0 to 13.6 +/- 0.62% after 10 minutes of treatment, and 8.1 +/- 1.24% after 30 minutes of treatment). The length of the remaining *Ttbk2^sca11/gt^* cilia reduced over time in a manner that was proportional to the other genotypes: for *Ttbk2^sca11/gt^* cells, cilia length at 30 post nocodazole was 63.2% of the starting length, compared with 65.4% for *Ttbk2^null/gt^* and 55.9% for WT (**Fig. 8D**). These data suggest that cilium stability is more compromised in *Ttbk2^sca11/gt^* cells than in *Ttbk2^null/gt^*, with cilia in *Ttbk2^sca11/gt^* cells rapidly lost in the presence of nocodazole, consistent with the dominant negative nature of the *sca11* allele.

To further investigate the role of TTBK2 in cilium stability and ciliary structure, we performed transmission electron microscopy (TEM) on neural tube sections from E10.5 embryos of each genotype to assess the cilia (**Fig. 8E-H**). We did not observe dramatic differences in the overall structure of cilia between *Ttbk2^gt/gt^* or *Ttbk2^null/gt^* and *Ttbk2^gt/+^*. By contrast, the structure of the cilia in *Ttbk2^sca11/gt^* embryos differs noticeably from the other genotypes in a number of ways. Consistent with our previous findings from *Ttbk2^null/null^* cells which have normal distal and subdistal appendages, we observe these structures in *Ttbk2^sca11/gt^* cilia, as well as the extension of axonemes that contain microtubules. However the microtubules appear less distinct than in the cilia of the other genotypes, with the proximal cilium/transition zone in particular having a less organized appearance. In addition, we frequently observed what appear to be vesicles within the ciliary axonemes of *Ttbk2^sca11/gt^* embryos, but not in cilia in embryos of the other genotypes. These TEM images, together with our data showing that polyglutamylated tubulin is reduced in Ttbk2 hypomorphic mutants, and in particular in *Ttbk2^sca11/gt^* mutants, suggest that TTBK2 is important for the structure and stability of the microtubule axoneme.

To examine the possible molecular mechanisms by which reduced TTBK2 could affect ciliary structure and stability, we tested whether a pathway important in cilium suppression and disassembly was altered upon reduced TTBK2 function. KIF2A is an atypical kinesin of the Kinesin 13 family that mediates microtubule depolymerization in a number of cellular contexts[31]. KIF2A was recently identified as a substrate of TTBK2 at the plus ends of cytoplasmic microtubules; in this context, phosphorylation of KIF2A by TTBK2 at S135 reduced the ability of KIF2A to bind microtubules, thereby impairing its depolymerase activity and stabilizing microtubules[10]. Given this association, we tested whether the localization of KIF2A was altered in *Ttbk2* mutant cells. In WT MEFs, KIF2A was localized to the centrosome and was also occasionally seen within the proximal ciliary axoneme (**Fig 9A**). Centrosomal localization was maintained in the *Ttbk2* mutant alleles. However, quantification of KIF2A intensity at the base of ciliated cells revealed that the level of KIF2A at the centrosome was increased in *Ttbk2^null/gt^* (mean pixel intensity of 43.1 +/- 1.96 A.U.) cells relative to *Ttbk^gt/+^* (mean pixel intensity of 19.98 +/- 0.92) or *Ttbk2^gt/gt^* (mean pixel intensity of 22.8 +/- 0.90 A.U). The intensity of KIF2A was further increased at the ciliary base of *Ttbk2^sca11/gt^* relative to all other genotypes (mean pixel intensity of 53.0 +/- 1.87 A.U., **Fig 9B**; p<0.0001). This suggests that KIF2A accumulates at the ciliary base when TTBK2 levels are reduced, where it could contribute either to structural defects in cilia, or to the observed reduction in ciliated cells by promoting cilium disassembly, or both.

**Figure 9.**
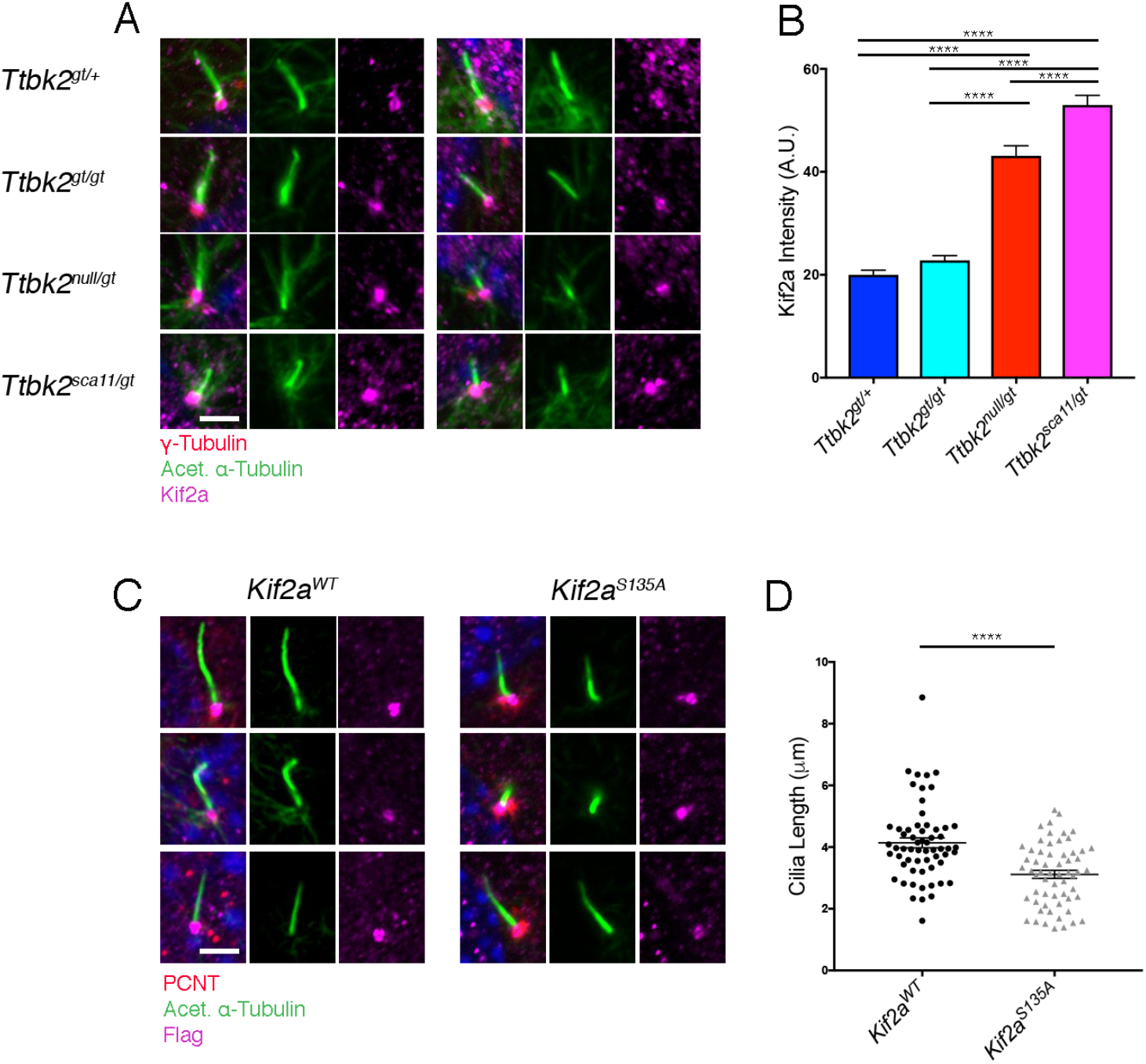
Kif2a accumulation at the base of *Ttbk2^sca11/gt^* cilia causes instability. **(A,B)** Immunostaining for KIF2A (magenta) in MEFs derived from embryos of the indicated genotypes of the Ttbk2 allelic series shown in (A). MEFs were serum starved for 48 hours. Centrosomes are labeled with γ-Tubulin (red) and cilia are stained for Acetylated α-Tubulin (green). Pixel intensity of KIF2A at the centrosome is quantified in (B). While we detected no significant difference between *Ttbk2^gt/+^* and *Ttbk2^gt/gt^*, *Ttbk2^null/gt^* and *Ttbk2^sca11/gt^* cells have increased levels of KIF2A at the centrosome, and KIF2A is further increased in *Ttbk2^sca11/gt^*. (1- way ANOVA, p<0.0001 for all). n=50 cilia pooled from 3 biological replicates. Scale bar = 5μm **(C,D)** *Ttbk2^gt/+^* cells were stably expressing either KIF2A^WT^ or a phospho-mutant variant of KIF2A, KIF2A^S135A^. Immunostaining is shown for the Kif2a construct (magenta), centrosomes stained for PCNT (red), and the ciliary axoneme labeled with Acetylated α-Tubulin (green). Three representative cilia are shown per condition. Cilia were shorter in *Ttbk2^gt/+^* cells when expressing KIFT2A^S135A^. Cilia length was determined and quantified in (D) (1-way ANOVA p<0.0001). Error bars represent SEM.

To assess whether and how loss or reduction in S135 phosphorylation of KIF2A may contribute to the ciliary phenotypes seen in the TTBK2 hypomorphic mutants, we tested the effects of expressing a non-phosphorylatable variant of KIF2A (S135A) in WT MEFs. We found that in MEFs over-expressing KIF2A^S135A^, cilia were significantly shortened compared to those overexpressing WT KIF2A (**Fig 9C, D**), although we did not observe a significant change in the percentage of ciliated cells between these conditions. Thus, we propose that increased activity of KIF2A in *Ttbk2* hypomorphic mutants contributes to defects in ciliary stability and structure, and to the exacerbated phenotypes observed in the *Ttbk2^sca11/gt^* embryos and neonates.

## Discussion

In this study, we show that the human SCA11-associated mutations to *Ttbk2* produce truncated proteins that interfere with the function of full-length TTBK2 in cilia formation. Consistent with our previous data showing that familial SCA11-associated mutations are unable to restore primary cilia in null mutant cells, our analysis of *Ttbk2^sca11/sca11^* mutants revealed a phenotype that is essentially indistinguishable from that of our previously described ENU-induced null allele. Like *Ttbk2^null/null^*, homozygous SCA11 mutants lack cilia in all tissues examined at E10.5, and the cells of these mutants exhibit an identical set of cellular defects to those of embryos lacking *Ttbk2*. These results indicate that TTBK2^SCA11^ truncations are completely unable to function in mediating ciliogenesis, despite having an intact kinase domain and producing a protein product[14]. This inability to function in ciliogenesis is likely the result of the SCA11-associated truncations lack of the C-terminus, which we and others have shown is required to target TTBK2 to the basal body and for its interaction with the distal appendage protein CEP164 [4,9,32].

In our prior studies we also found that expression of TTBK2^SCA11^-GFP in WT fibroblasts led to a modest but significant reduction in ciliogenesis [4], consistent with the classical definition of a dominant negative [33]. We hypothesized based on this that the SCA11-associated mutations to *Ttbk2* function as antimorphic alleles. In the current work, we present two major lines of evidence in support this hypothesis. First, we found that expression of WT TTBK2-GFP only partially rescues cilia formation in *Ttbk2^sca11/sca11^* mutant cells whereas full rescue is achieved by stable expression of the same construct in *Ttbk2^null/null^* cells. This is seen both at the level of ciliogenesis, where many fewer ciliated cells are found in rescued *Ttbk2^sca11/sca11^* cells and also with respect to the structure of the cilium: ARL13B localization is significantly impaired in the rescued *Ttbk2^sca11/sca11^* relative to rescued null mutant cells. Second, we also present genetic evidence for the dominant negative function of *Ttbk2^sca11^*. The combination of *Ttbk2^sca11^* with a hypomorphic allele that produces a reduced amount of TTBK2 protein (*Ttbk2^gt^*) results in more severe phenotypes than the null allele in combination with *Ttbk2^gt^* on the same genetic background.

We propose a model wherein TTBK2’s functions in cilium assembly are highly dosage sensitive, with alterations in the amount of functional TTBK2 protein below a certain threshold causing a range of phenotypes related to defects in ciliary trafficking and signaling. In human SCA11 patients, the presence of SCA11 truncated protein is sufficient to cause a phenotype limited to a specific tissue- the cerebellum. In mice, we did not identify any changes in the architecture of the cerebellum between *Ttbk2^sca11/+^* animals and their WT siblings by 3 months of age. While we can’t yet exclude the emergence of more subtle defects occurring at advanced age, it does not appear one allele of *Ttbk2^sca11^* is sufficient to cause phenotypes recapitulating human SCA11 in the presence of a second WT allele of *Ttbk2*, (ie *Ttbk2^sca11/+^*) in mice. However, on a sensitized background with a reduced amount of full-length TTBK2, the dominant negative effects of TTBK2^SCA11^ become apparent, such as in the allelic series.

Our studies of the ciliary defects occurring in the *Ttbk2* allelic series have also yielded valuable insights about the role of TTBK2 in cilia formation and trafficking. Our prior work based on a null allele of *Ttbk2* demonstrated the essential role played by this kinase in initiating cilium assembly upstream of IFT. However, examination of hypomorphic alleles in this study points to additional requirements for TTBK2 following initial cilium assembly. For example, cilia are shorter in cells derived from all of the hypomorphic *Ttbk2* alleles compared with WT or *Ttbk2^gt/+^* cells, pointing to a role for TTBK2 in cilia structure and trafficking. Identifying the molecular targets of TTBK2 in both cilium initiation and in ciliary trafficking and/or stability will be critically important to our understanding of the pathways that regulate ciliogenesis.

We identified modest disruptions in the concentration of IFTB and IFTA components in our *Ttbk2* hypomorphic allelic combinations, consistent with a role for TTBK2 particularly in maintaining the basal body pools of IFT proteins, in addition to the requirement for TTBK2 in the recruitment of IFT components to the basal body that we identified previously in our analysis of *Ttbk2* null mutant cells[4]. Identifying the mechanisms by which TTBK2 mediates the localization of IFT proteins to the basal body, as well as testing whether TTBK2 contributes to IFT mediated ciliary trafficking will be a focus of our future studies.

We have also uncovered a role for TTBK2 in maintaining the stability of the ciliary axoneme, with these defects becoming particularly evident in *Ttbk2^sca11/gt^* mutant cells. Consistent with a requirement for TTBK2 in cilia structure, KIF7 is reduced in *Ttbk2^sca11/gt^* cells compared with *Ttbk2^null/gt^* cells with respect to the percentage of cilia that are positive for KIF7. The *Ttbk2^sca11/gt^* cells also exhibit a subset of the defects found in *Kif7^-/-^* cells, including a reduction in polyglutamylated tubulin[25]. Unlike *Kif7^-/-^* cells however, we did not observe any reduction in tubulin acetylation in any of the *Ttbk2* hypomorphic cells. The *Ttbk2^sca11/gt^* cells do exhibit increased instability in the presence of nocodazole. The highly modified microtubules of the cilium are typically relatively resistant to this microtubule-depolymerizing drug [25,34], and in WT or *Ttbk2^null/gt^* cells neither the proportion of ciliated cells nor the length of the cilium changes dramatically when the cells are treated with nocodazole for up to 30 minutes. In contrast, in the *Ttbk2^sca11/gt^* cells the percentage of ciliated cells drops dramatically upon treatment with nocodazole, consistent with a requirement for TTBK2 in the stability of the axonemal microtubules. In addition, increased levels of the microtubule depolymerizing kinesin KIF2A are present at the centrosome of ciliated cells in the *Ttbk2* hypomorphic mutants, with the highest amounts seen in *Ttbk2^sca11/gt^* cells. This suggests that TTBK2 may oppose the activity the PLK1-KIF2A cilium disassembly pathway, and that an increase in the activity of this pathway in the *Ttbk2* hypomorphic mutants contributes to the reduction in ciliated cells, in addition to defects in cilium stability.

Loss of tubulin glutamylation in cilia has also recently been shown to perturb ciliary trafficking and the enrichment of HH pathway components within cilia upon stimulation of cells with SAG [35]. Thus, the reduction in SMO enrichment that we observed in the *Ttbk2^sca11/gt^* cells could result from the additional reduction in tubulin glutamylation we see within the cilia of these cells relative to other genotypes, with disrupted trafficking in HH pathway components in turn leading to exacerbated embryonic phenotypes related to HH-dependent patterning.

The additional impairment of TTBK2 function in the *Ttbk2^sca11/gt^* animals results in a greater perturbation of cilia than the defects seen in *Ttbk2^null/gt^* cells. These include reduced numbers of cilia, disrupted cilium stability, and impaired trafficking of signaling molecules such as SMO to the axoneme, although we have not yet precisely defined the biochemical mechanisms by which the human disease-associated truncations interfere with TTBK2 function. Our data argue against a model where TTBK2^SCA11^ directly binds to full length TTBK2 and inhibits its function through a direct association. Rather, it seems more likely that TTBK2^SCA11^, having lost critical regulatory motifs as well as the ability to efficiently translocate to the centrosome, may sequester some important TTBK2 substrate or substrates, resulting in the further impairment of cilia structure and signaling that in turn causes the modest exacerbations in SHH-dependent developmental patterning.

While our data indicate that the SCA11-associated *Ttbk2* mutations interfere with cilia formation and stability, pointing to a strong possibility that SCA11 pathology is related to disrupted ciliary signaling, we cannot exclude the possibility that TTBK2 has non-ciliary roles within the brain that could also contribute to neural degeneration. For example, TTBK2 phosphorylates Synaptic Vesicle Protein 2A, and this event is important for the formation and release of synaptic vesicles [36]. The mechanisms of TTBK2 regulation and the specific substrates of this kinase in cilium assembly, as well as possible non-ciliary roles for TTBK2 within the brain are key topics in our ongoing research. Having shown that TTBK2^SCA11^ is both unable to mediate cilium assembly and also impairs the function of TTBK2^WT^ in ciliogenesis, another important area of investigation is the relationship between cilia and ciliary signaling pathways and the maintenance of neural connectivity and function.

## Materials and Methods

### Ethics statement

The use and care of mice as described in this study was approved by the Institutional Animal Care and Use Committees of Memorial Sloan Kettering Cancer Center (approval number 02-06-013) and Duke University (approval numbers A246-14-10 and A218-17-09). Euthanasia for the purpose of harvesting embryos was performed by cervical dislocation, and all animal studies were performed in compliance with internationally accepted standards.

### Mouse Strains

We used two previously described alleles of *Ttbk2*: *Ttbk2^null^* is an ENU-induced allele (also called *Ttbk2^bby^*) [4], and *Ttbk2^sca11^* is a knockin recapitulating one of the familial SCA11-associated mutations [14]. Genotyping for both of these alleles was performed as previously described. *Ttbk2* “knockout first” genetrap (*Ttbk2^tm1a(EUCOMM)Hmgu^*, here referred to *Ttbk2^gt^*) targeted ES cells were purchased from the European Mutant Mouse Consortium. One clone (HEPD0767_5_E08, parental ESC line JM8A3.N1, agouti) was injected into host blastocysts by the Mouse Genetics Core Facility at Sloan Kettering Institute. Resulting chimeric male mice were bred to C57BL/6 females to test germline transmission and obtain heterozygous mice. PCR genotyping (F: ATACGGTTGAGATTCTTCTCCA, R1: TCTAGAGAATAGGAACTTCGG, R2: TGCAATTGCATGACCACGTAGT) yields a band corresponding to the mutant allele at 407bp and to the WT allele at 762bp.

### Embryo and tissue dissection

To obtain embryos at the identified stages, timed matings were performed with the date of the vaginal plug considered embryonic day (E) 0.5. Pregnant dams were sacrificed by cervical dislocation and embryos were fixed in either 2% (E11.5 or earlier) or 4% (later than E11.5) paraformaldehyde (PFA) overnight at 4C. For cryosectioning, tissue was cryoprotected in 30% Sucrose overnight and embedded in Tissue Freezing Medium (General Data TFM-5). Tissue was sectioned at 16μm thickness.

To harvest tissues from adult mice, animals were anesthetized with 12.5mg/mL Avertin, and a transcardially perfused with of Phosphate Buffered Saline (PBS) followed by 4% PFA. Kidneys and brains were dissected and incubated in 4% PFA for an additional 2 hours at 4°C. Tissue was then prepared for cryosectioning as described above.

### Cell culture and immunostaining

MEFs were isolated from embryos at either E10.5 or E12.5, and maintained as previously described [37]. To induce cilia formation, cells were shifted from 10% to 0.5% fetal bovine serum (FBS) and maintained in low serum conditions for 48 hours. Cells were grown on coverslips and fixed in 4% Paraformaldehyde (PFA) in Phosphate Buffered Saline (PBS) for 5 minutes at room temperature followed by methanol for 5 minutes at −20C. Cells were then washed in PBS + 0.2% Triton X-100 (PBT) and blocked in PBT + 5% FBS + 1% bovine serum albumin for 30 minutes. Cells were then incubated with primary antibodies diluted in blocking solution overnight at 4 °C, and finally incubated with Alexa-coupled secondary antibodies and DAPI in blocking solution for 30 minutes at room temperature and affixed to slides for microscopy. Embryonic and adult tissue sections were collected onto slides, dried, washed in PBT + 1% serum, and incubated with primary antibodies as described above.

### Producing Kif2a phospho-mutant cell lines

The human Kif2a Gateway-ready clone was obtained from the Human ORFeome Collection (Dharmacon clone BC031929). The KIF2A^S135A^ mutation was introduced via site directed mutagenesis using the Quick Change II Mutagenesis Kit (Agilent). Using Gateway LR Clonase (Invitrogen), both KIF2A^WT^ and KIF2A^S135A^ clones were transferred into Gateway Destination vectors compatible for retroviral vector expression modified to contain eGFP and FLAG tags. Retroviral transduction was carried out as previously reported [4].

### Antibodies

The SMO antibody was raised in rabbits (Pocono Rabbit Farm and Laboratory Inc.) using antigens and procedures described[20]; diluted 1:500. Antibodies against KIF7 [23] (1:1000), ARL13B [35] (1:2000), GLI2 [36] (1:2000) and TTBK2 [14] have been previously described. Commercially available antibodies used in these studies were: mouse anti-NKX2.2, ISL1 (Developmental Studies Hybridoma Bank, each 1:10); mouse anti-Pericentrin, (BD Biosciences #611814, 1:500) γ-Tubulin (Sigma SAB4600239, 1:1000), Acetylated α-Tubulin (Sigma T6793, 1:1000), polyglutamylated Tubulin (Adipogen AG-20B-0020, 1:2000); rabbit anti-IFT88 (Proteintech 13967-1-AP, 1:500), rabbit anti-IFT81 (Proteintech 1174-1-AP, 1:1000), rabbit anti-IFT140 (Proteintech 17460-1-AP 1:500), mouse anti-FLAG (Sigma F1804, 1:1000), rabbit anti-KIF2A (Abcam ab37005, 1:500), TTBK2 (Proteintech 15072-1-AP, 1:1000), Calbindin (Cell Signaling Technology 13176-S, 1:250), VGLUT2 (EMD Millipore AB2251, 1:2500).

### Microscopy

Immuno-fluorescence images were obtained using a Zeiss AxioObserver wide field microscope equipped with an Axiocam 506mono camera and Apotome.2 optical sectioning with structured illumination. Z-stacks were taken at 0.24μm intervals. Whole mount images of embryos and tissues were captured with a Zeiss Discovery V12 SteREO microscope equipped with an Axiocam ICc5 camera. Image processing and quantifications were performed using ImageJ. To quantify the signal intensities of ciliary proteins, Z stack images were captured using the 63X objective. A maximum intensity projection was then created for each image using ImageJ, background was subtracted. Cilia were identified by staining with Acetylated α-Tubulin and γ- Tubulin. Each cilium or portion of the cilium was highlighted using either the polygon tool or the line tool (for line-scan analysis), and the mean intensity was recorded for the desired channel (measured on an 8 bit scale), as described [25]. To measure the mean intensity, ImageJ software was used to calculate total intensity divided by the area selected. Measurements taken within the cilium therefore take into account the length per measurement recorded. Statistical analysis was done with the Prism7 statistical package (GraphPad).

### Western blotting and immunoprecipitation

HEK-293T cells were transfected with constructs for tagged proteins of interest using Lipofectamine 3000 (Thermo Fisher) according to the manufacturer’s instructions. Constructs used were TTBK2^FL^-GFP, TTBK2^FL^-V5, TTBK2^SCA11^-V5 (1-443aa, Ttbk2^Cterm^-GFP (306-1243aa).

For western blots, cells or tissues were lysed in buffer containing 10mM Tris/Cl pH7.5, 150mM NaCl, 0.5mM EDTA, 1% Triton, 1mM protease inhibitors (Sigma #11836170001) and 25mM β- glycerol phosphate (Sigma 50020), and total protein concentration was determined using a BSA Protein Assay Kit (Thermo Fisher #23227).

For co-IP experiments, cells were lysed in buffer containing 20mM Tris-HCl pH7.9, 150mM NaCl, 5mM EDTA, 1% NP-40, 5% glycerol, 1mM protease inhibitors and 25mM β-glycerol phosphate. Immunoprecipitation of lysates was performed using analysis was done using GFP-Trap beads (Chromotek GTA-20) blocked with 3% BSA in Co-IP lysis buffer overnight prior to pull-down. rabbit α-GFP (Invitrogen A11122, 1:10,000), mouse α-V5 (Invitrogen R96025, 1:7,000), HRP-conjugated secondaries (Jackson ImmunoResearch).

### Cerebellum Quantification

Quantification of the cerebellar tissue was done using ImageJ software. Images for the molecular layer analysis were taken at 20x. For measuring the molecular layer, a line was drawn from the top of the PC cell soma to the pial surface and the distance was recorded. That same line was then brought down from the pial surface to the top of the nearest VGLUT2 puncta along that line, the distance was recorded, and a ratio was calculated. Measurements were pooled equally from both sides of the primary folia of the cerebellum, and from four slices per animal. Images for the VGLUT2 analysis were 10μm thick z-stacks taken at 63x. VGLUT2 puncta analysis was performed using the ImageJ “Analyze Particles” plug-in with the following stipulations: Size exclusion: 0.5-infinity, Circularity: 0-1. Measurements were pooled from 5 areas in the cerebellum, and from four slices per animal.

### RT-PCR

RNA was extracted from brains dissected from p30 animals using the Qiagen RNeasy Mini Kit (Qiagen, 74104). cDNA was then made from 1μg of RNA using the BioRad iScript cDNA Synthesis Kit (BioRad, 1708891). PCR primers were designed to span the exon 4-5 boundary of *Ttbk2* (F: ATGCTCACCAGGGAGAATGT, R: TGCATGACCACGTAGTTGAAA), lacZ (F: AGCAGCAGTTTTTCCAGTTC, R: CGTACTGTGAGCCAGAGTTG), and GAPDH (F: ACCACAGTCCATGCCATCAC, R: TCCACCACCCTGTTGCTGTA).

### Statistics

Indicated statistical comparisons were performed using Graphpad Prism7. For multiple comparisons, a Tukey-Kramer post-hoc test was performed.

## Acknowledgements

We are grateful to Jonathan Eggenschwiler and Dario Alessi for reagents. We thank Don Fox, Debby Silver, and Karel Liem for their comments on the manuscript.

## Supplementary Information

**Figure S1.**
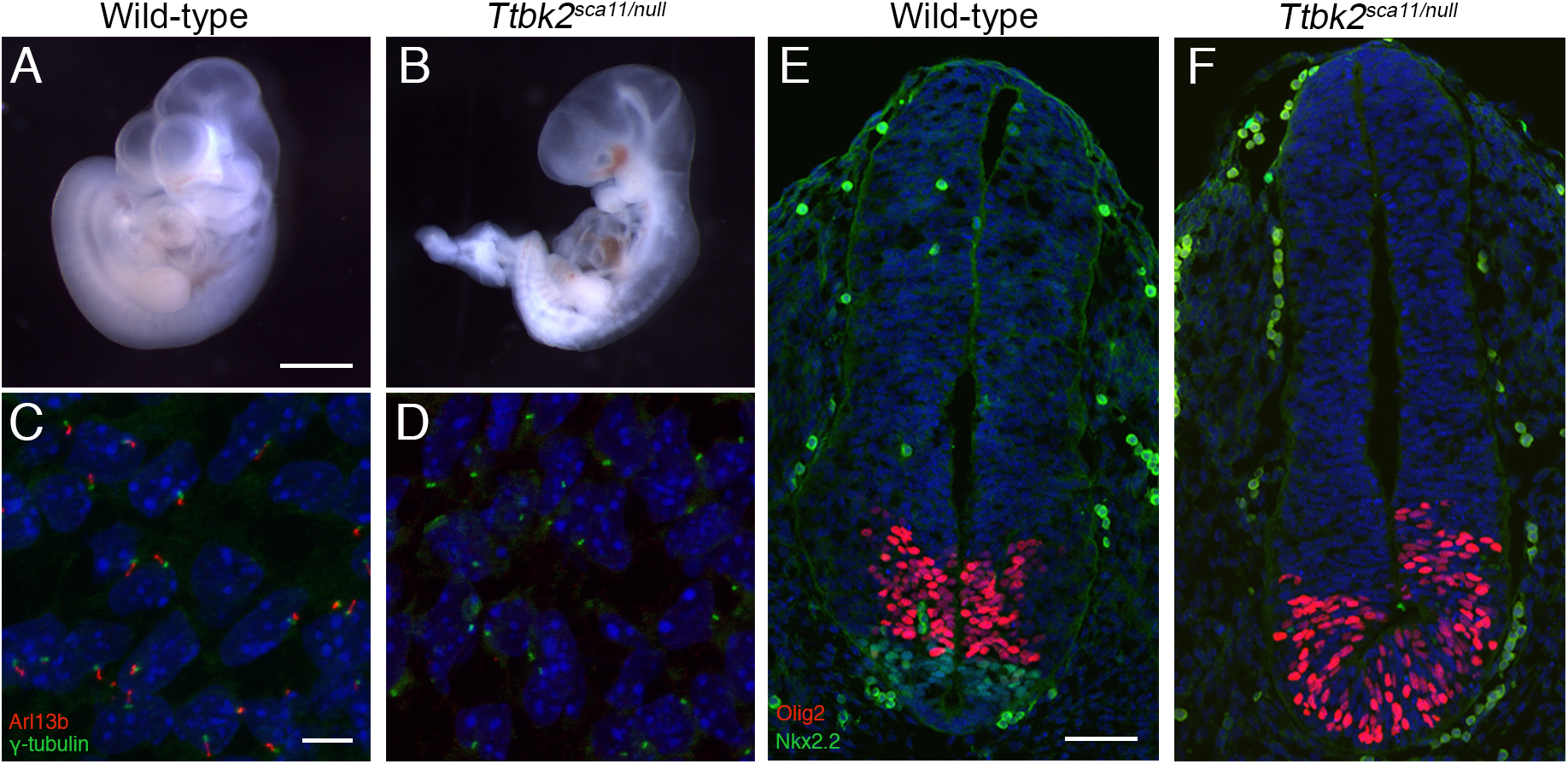
*Ttbk2^sca11/null^* embryos phenocopy *Ttbk2^null/null^* and *Ttbk2^sca11/sca11^* embryos. **(A,B)**Representative Wild type (A) and Ttbk2^*sca11/null*^ (B) E10.5 embryos. Scale bar = 1mm. **(C,D)** Mesenchymal cells surrounding neural tube of E10.5 Wild-type (C) and *Ttbk2^sca11/null^* (D) embryos. Sections are immunostained for cilia using ARL13b (red) and γ-Tubulin (green). Scale bar = 20μm. **(E,F)**Transverse sections of E10.5 neural tubes of Wild-type (E) and Ttbk2^*sca11/null*^ (F). Sections are immunostained for NKX2.2 to label V3 interneuron progenitors and OLIG2 to label motor neuron progenitors. Scale bar =100μm.

**Figure S2.**
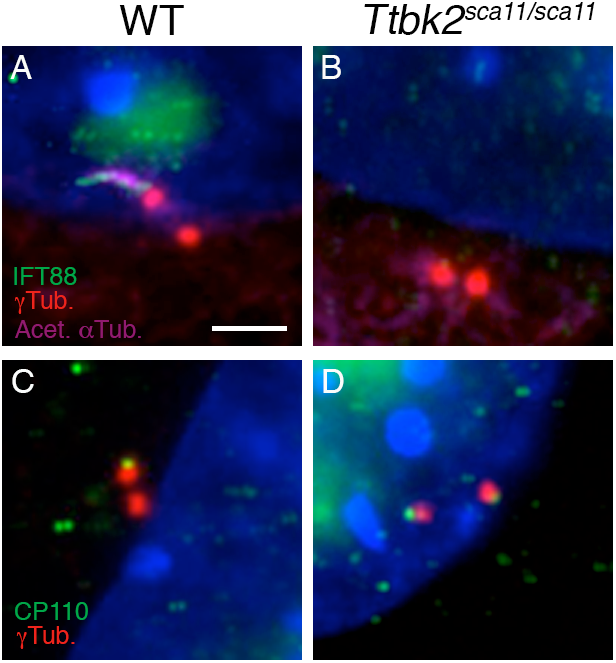
Cellular defects in *Ttbk2^sca11/sca11^* cells recapitulate those seen in *Ttbk2^null/null^*. **(A,B)** MEFs of the indicated genotype were serum starved for 48 hours and immunostained for IFT88 (green) as well as γ-Tubulin (red) to label centrosomes and Acetylated α-Tubulin (magenta) to label the axonemes of cilia. *Ttbk2^sca11/sca11^* cells lack cilia and also lack IFT88 at the mother centriole. **(C,D)**. Serum starved MEFs were treated as above and stained for CP110 (green) and γ- Tubulin (red). *Ttbk2^sca11/sca11^* cells retain CP110 on both centrosomes in the absence of serum. Scale bar = 5μm.

**Figure S3.**
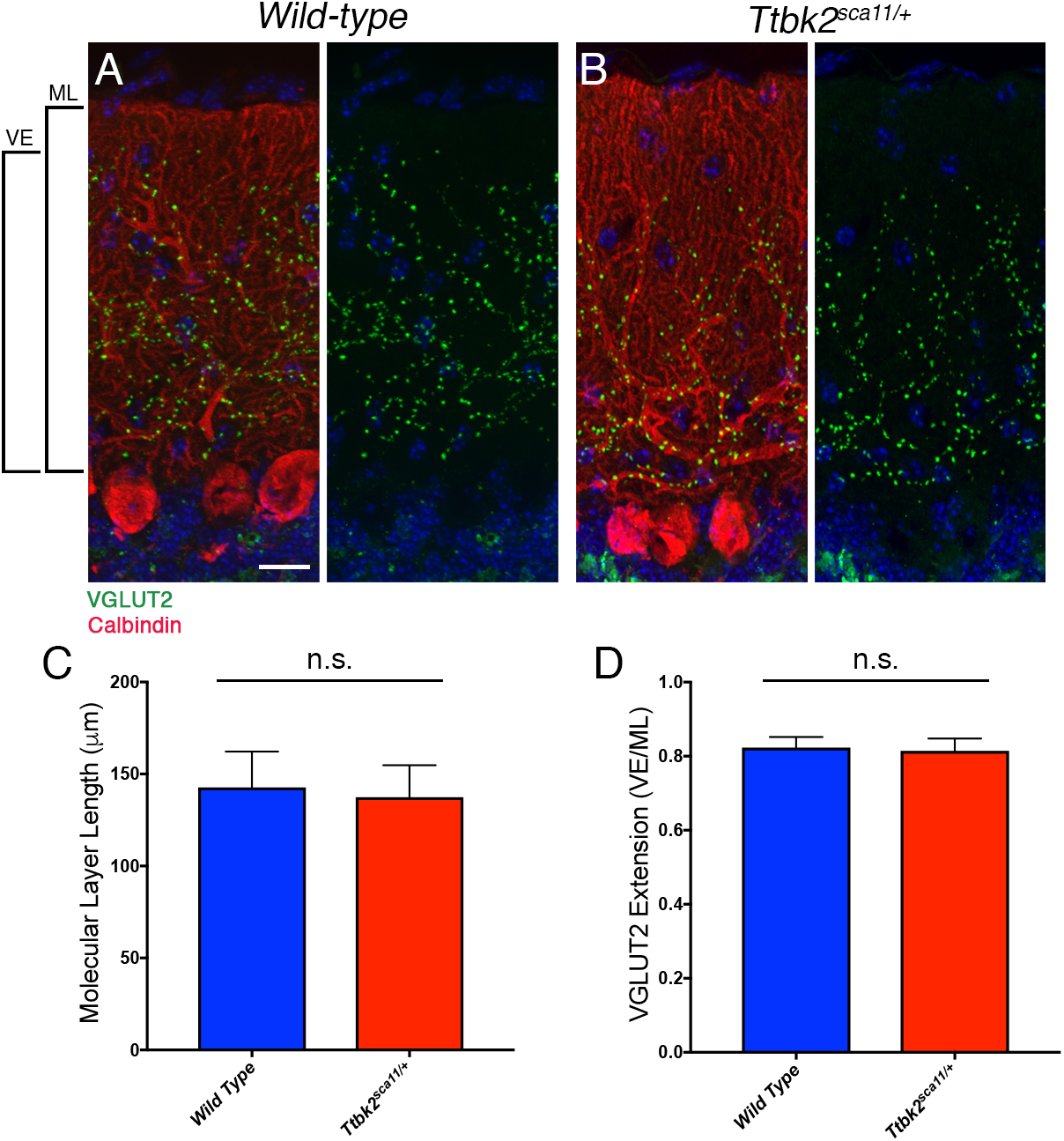
*Ttbk2^sca11/+^* mice do not show signs of cerebellar degeneration. **(A,B)** Representative sagittal sections through the cerebellum of mice of the indicated genotype. Purkinje cells are labeled with Calbindin (red) and excitatory synapses from the climbing fibers onto the Purkinje cell dendrites are labeled with VGLUT2 (green). Scale bar = 50μm. **(C,D)** Measurements for the molecular layer thickness and VGLUT2 extension were made from four separate primary folia, pooled from three individual mice for each condition. Error bars denote SEM. The molecular layer thickness and VGLUT2 extension are unchanged between wild-type and *Ttbk2^sca11/+^*.

**Figure S4.**
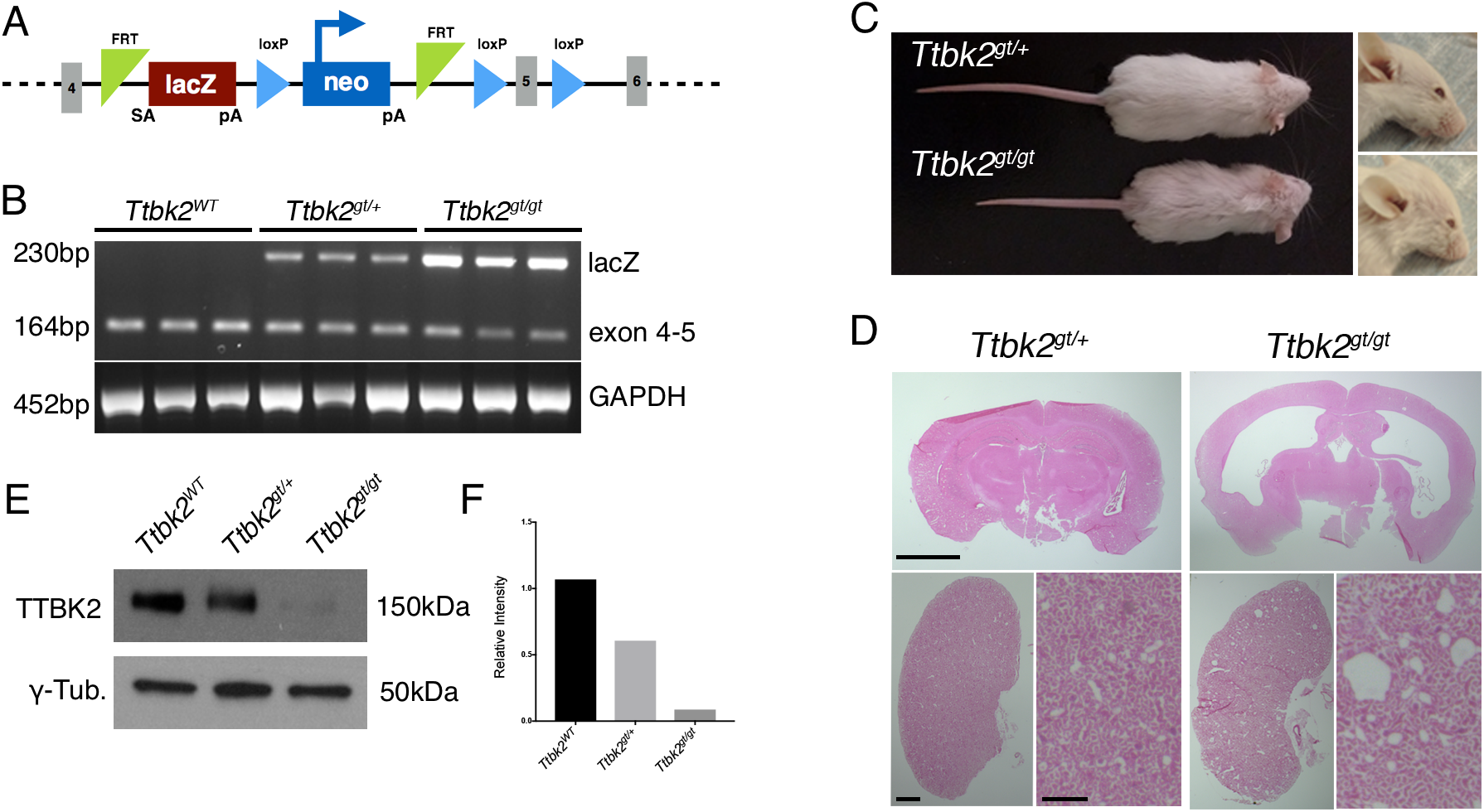
*Ttbk2^gt^* is a hypomorphic allele of *Ttbk2*. **(A)** Schematic of the *Ttbk2* gene trap (*Ttbk2^gt^*) targeting design (schematic adapted from International Mouse Phenotyping Consortium). **(B)** RT-PCR analysis of WT splicing in *Ttbk2^gt/gt^*. RNA from 3 biological replicate brains per genotype was used, and primers targeting the exon 4-5 boundary show that WT transcript is still produced in *Ttbk2^gt/gt^* mice. **(C)** P30 mice showing phenotypic differences between *Ttbk2^gt/+^* and *Ttbk2^gt/gt^*. **(D)** H&E staining of neural cortex and kidney tissue from 6mo old *Ttbk2^gt/gt^* mice showing hydrocephaly and polycyctic kidneys. Scale bar = 1mm. **(E)** Western blot showing decreased TTBK2 protein levels (150kDa) in lysates from *Ttbk2^WT^*, *Ttbk2^gt/+^*, and *Ttbk2^gt/gt^* brains. γ-Tubulin is a loading control. **(F)** Quantification of the relative intensity of the Western blot bands for TTBK2. TTBK2 protein levels in *Ttbk2^gt/gt^* brain lysate is about 8.2% of the amount in *Ttbk2^WT^* brain lysate.

**Figure S5.**
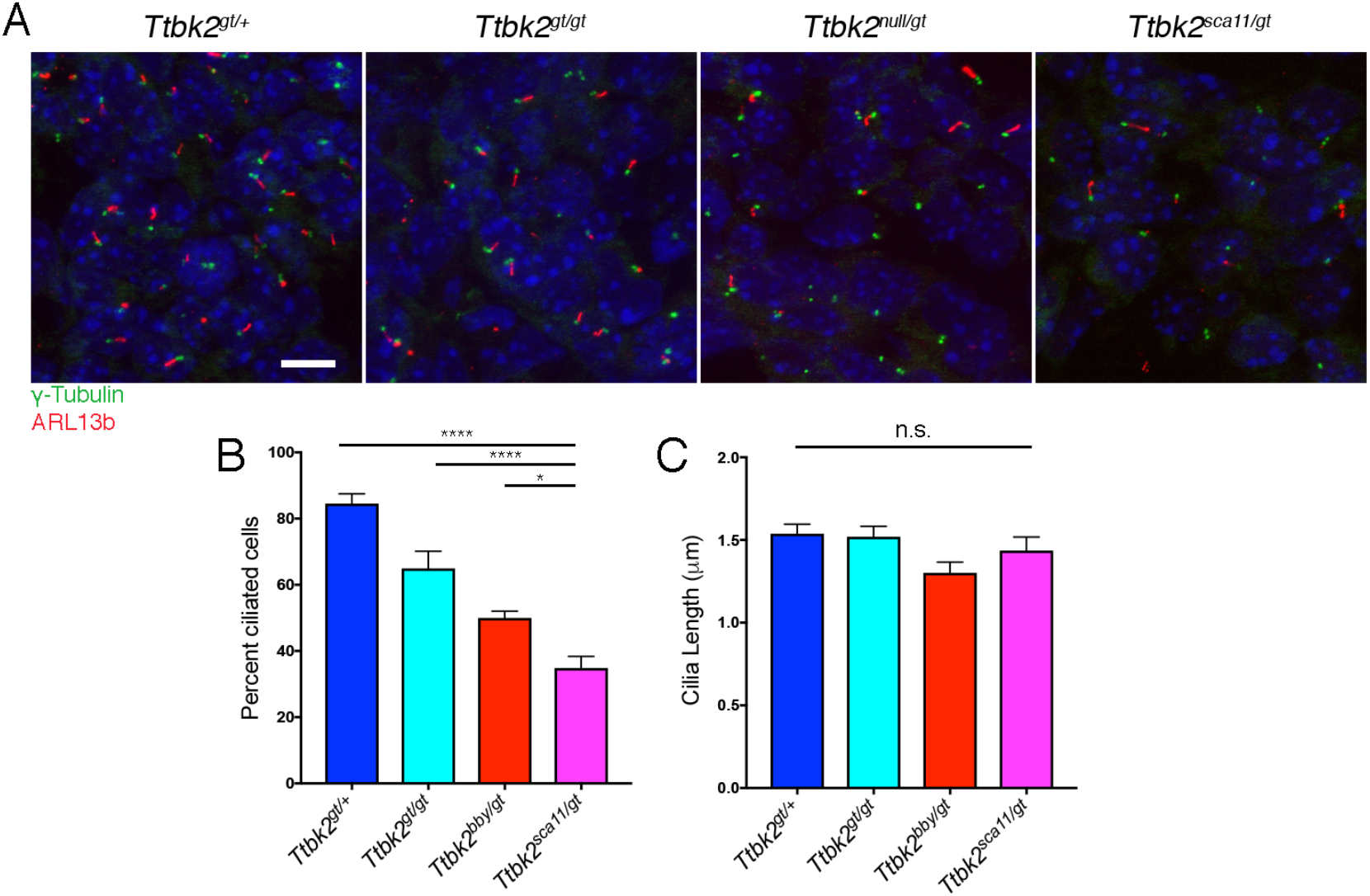
Cilia are less abundant in *Ttbk2* allelic series *in vivo.* **(A)** Representative images of mesenchymal cells surrounding the neural tube of E10.5 embryos of the indicated genotype. Cilia were immunostained for ARL13b (red) to label cilia and γ- Tubulin (green) to label centrosomes. Scale bar = 20μm. **(B)** Quantification of the percentage of ciliated cells in the mesenchyme of the indicated genotype. Cilia are less abundant in *Ttbk2^sca11/gt^* embryos. Statistical comparison was performed by 1-way ANOVA with Tukey-Kramer post-hoc test. (p<0.0001 vs *Ttbk2^gt/+^*; p<0.0001 vs *Ttbk2^gt/gt^*; p=0.0279 vs *Ttbk2^null/gt^*). n=two fields of view, three biological replicates, over 1000 total cells counted per genotype. **(C)** Quantification of cilia length in the mesenchyme of the indicated genotype. Cilia length is not statistically significantly different across the *Ttbk2* allelic series in the embryonic mesenchyme. n=50 cilia pooled from 3 biological replicates.

**Table S1.** Numbers of mutants obtained from allelic series crosses.

| Cross | Stage | Total Number | Number of mutants | Percent Mutant |
| --- | --- | --- | --- | --- |
| Null X Null | E9.5-E10.5 | 330 | 63 | 19.1 |
| Sca11 X Sca11 | E9.5-E10.5 | 94 | 20 | 21.3 |
| GT X GT | E12.5-E14.5 | 81 | 17 | 20.9 |
|  | E17.5-P0 | 183 | 44 | 24.0 |
| GT X Null | E12.5-E14.5 | 67 | 21 | 31.3 |
|  | E17.5-P0 | 43 | 8 | 18.6 |
| GT X SCA11 | E12.5-E14.5 | 99 | 22 | 22.2 |
|  | E17.5-P0 | 63 | 6 | 9.5 |

**Table S2.** Summary of gross phenotypes observed from allelic series crosses.

| Cross | Stage | Number of mutants | Number polydactyly | Number holopros. | Number midbrain defect | Number forebrain defect |
| --- | --- | --- | --- | --- | --- | --- |
| Null X Null | E9.5-E10.5 | 63 | N/A | 63 | 63 | N/A |
| Sca11 X Sca11 | E9.5-E10.5 | 20 | N/A | 20 | 20 | N/A |
| GT X GT | E12.5+ | 61 | 4 | N/A | N/A | 0 |
| GT X Null | E9.5-E10.5 | 10 | N/A | 0 | 0 | N/A |
|  | E12.5+ | 29 | 29 | N/A | N/A | 19 |
| GT X SCA11 | E9.5-E10.5 | 12 | N/A | 4 | 4 | N/A |
|  | E12.5+ | 28 | 28 | N/A | N/A | 22 |

